# Penalized likelihood optimization for censored missing value imputation in proteomics

**DOI:** 10.1101/2023.11.09.566355

**Authors:** Lucas Etourneau, Laura Fancello, Samuel Wieczorek, Nelle Varoquaux, Thomas Burger

**Affiliations:** Univ. Grenoble Alpes, CNRS, CEA, Inserm, BGE UA13, ProFI FR2048, 38000, Grenoble, France; TIMC, Univ. Grenoble Alpes, CNRS, Grenoble INP, 38000, Grenoble, France

**Keywords:** (1) Mass Spectrometry Based Proteomics, (2) Imputation of Missing Not At Random Values, (3) Multi-Omic Imputation, (4) Covariance Matrix Estimation, (5) Penalized Likelihood Maximization

## Abstract

Label-free bottom-up proteomics using mass spectrometry and liquid chromatography has long been established as one of the most popular high-throughput analysis workflows for proteome characterization. However, it produces data hindered by complex and heterogeneous missing values, which imputation has long remained problematic. To cope with this, we introduce Pirat, an algorithm that harnesses this challenge using an original likelihood maximization strategy. Notably, it models the instrument limit by learning a global censoring mechanism from the data available. Moreover, it estimates the covariance matrix between enzymatic cleavage products (*i*.*e*., peptides or precursor ions), while offering a natural way to integrate complementary transcriptomic information when multi-omic assays are available. Our benchmarking on several datasets covering a variety of experimental designs (number of samples, acquisition mode, missingness patterns, etc.) and using a variety of metrics (differential analysis ground truth or imputation errors) shows that Pirat outperforms all pre-existing imputation methods. Beyond the interest of Pirat as an imputation tool, these results pinpoint the need for a paradigm change in proteomics imputation, as most pre-existing strategies could be boosted by incorporating similar models to account for the instrument censorship or for the correlation structures, either grounded to the analytical pipeline or arising from a multi-omic approach.

## 1 Introduction

Bottom-up label-free LC-MS/MS (tandem mass spectrometry coupled with liquid chromatography) stands out as the most widely used method to characterize the proteome of biological samples. These assays typically yield an abundance matrix, where each row corresponds to a sample, each column a protein. Yet, moving from the massive amount of MS signals generated to such abundance matrix requires robust, mathematically well-grounded, and fast evolving software, posing computational and theoretical challenges. Notably two difficulties specific to bottom-up workflows arise. First, protein abundances are not directly measured: Sample preparation involves proteolysis (*i*.*e*., the cleavage of each protein into several *peptides* using an enzyme like trypsin) and MS measurements require the peptides to be ionized beforehand (to which we refer to as *precursor ions*, or simply *precursors*). Thus, inferring the identity and quantity of proteins requires two aggregation steps: from precursors to peptides, and from peptides to proteins. Second, raw LC-MS/MS data are hampered by a large number of missing values (MVs) in the abundance matrices: the overall MVs rate can reach 50% [Lazar et al., 2016]—whether that is at the level of precursor, peptide, or protein abundances; and at least 50% of peptides usually have at least one MV [Liu and Dongre, 2021].

The origin of MVs is complex and multi-faceted. Since the work of Karpievitch et al. [2012], it is customary to separate MVs into two types. The first one gathers all the MVs resulting from the various workflow imperfections (such as peptide ionization issues, enzymatic miscleavages, too complex samples, false peptide identifications, *etc*.) and which may affect proteins broadly randomly, regardless of their abundance. The second category corresponds to the censorship mechanism resulting from the instrument limit: precursors with an abundance below this limit yield MVs. A major issue with this censorship relates to its stochastic and dynamic nature, as the MS range changes along the acquisition process [Vidova and Spacil, 2017]. As a downside of the dramatic proportion and complex nature of MVs, analysing only fully observed biomolecules (be they precursors, peptides or proteins) is usually not considered, as it would result in discarding too much biologically relevant information. While it is theoretically safer to conduct the analysis without inferring MVs [Little and Rubin, 2019], *e*.*g*., identifying differentially expressed proteins using models that cope with MVs [Chen et al., 2014, Ryu et al., 2014, O’brien et al., 2018, Goeminne et al., 2020, Chion and Leroy, 2023], many downstream investigation techniques require in practice to impute first MVs (*i*.*e*., to estimate the missing abundances). Yet, despite extensive research, this remains a fundamental problem of computational proteomic research.

Drawing from a vast body of literature that spans a decade, some consensus on the subject has emerged. First, imputing missing protein abundances is suboptimal [Lazar et al., 2016] as a result of the effect of the peptide-to-protein aggregation on missing values. However, whether imputation should be performed at precursor- or peptide-level has not been investigated yet. Second, it is insightful to distinguish MVs according to the following well-acknowledged statistical categories [Little and Rubin, 2019]: Missing Completely At Random (MCAR), where the probability that a value is missing does not depend on any observed or missing value in the data; Missing At Random (MAR), where the same probability may only depend on observed values; and Missing Not At Random (MNAR) in any other case. Accordingly, in bottom-up label free proteomics, MVs resulting from the lower instrumental limit are classically assumed to follow an MNAR mechanism [Karpievitch et al., 2012, Webb-Robertson et al., 2015, Lazar et al., 2016], while other MVs, in absence of sufficiently refined MAR-based description, are modelled as MCARs. Third, the algorithms providing the best imputations are not the same depending on whether MVs are classified as MCARs or MNARs [Lazar et al., 2016]. This observation has motivated the development of meta-imputation tools, *i*.*e*., algorithms taking as input one or several imputation methods(s), and delivering as output a more refined imputation result *e*.*g*., [Wei et al., 2018, Ma et al., 2020, Giai Gianetto et al., 2020, Gardner and Freitas, 2021, Wang et al., 2022, Chion et al., 2022], as well as diagnosis tools capable of proposing an imputation strategy tailored to the data specificities [Wang et al., 2020, Kong et al., 2022]. Regardless of their practical interest, leveraging those approaches is only possible if classical imputation methods are available in the first place.

Numerous exhaustive reviews propose comparisons between a wide range of imputation algorithms, see for example [Lazar et al., 2016, Karpievitch et al., 2012, Webb-Robertson et al., 2015, Jin et al., 2021, Liu and Dongre, 2021, Shen et al., 2022], as well as Section 3.2. Although those works generally concur on the least accurate approaches, they do not agree on the most accurate ones, which depend on the dataset or on the experimental design. Moreover, while MCAR/MAR-devoted methods have established their robustness in many scientific domains where MVs have hardly specific behaviors, the MNAR-devoted methods used in proteomics are often simple and univariate, thus poorly informative [Chion and Leroy, 2023] and leading to larger imputation errors. A notable exception is msImpute [Hediyeh-zadeh et al., 2023], whose very recent publication has unveiled promising results. Contrarily to most anterior methods, msImpute simultaneously tackle MCARs and MNARs, by estimating the type of each MV and interpolating MAR and MNAR imputation distributions accordingly. Yet, some parameters (as the barycenter weights or the MNAR assumption) are fixed in a predetermined manner, which may hinder the generalization capabilities of the approach. Pushing the logic a step further, few works [Chen et al., 2014, Li and Smyth, 2023] propose to bypass any MCAR/MNAR distinction, assuming the relevant model should directly estimate the probability of missing depending on the abundance. Among them, PEMM [Chen et al., 2014] has been an important source of inspiration for this work. Considering an analytical approximation of a left-censoring mechanism, it uses the Expectation-Maximization algorithm to maximize a penalized likelihood model of the mean and covariance matrix of the protein-level data, yielding natural parametric imputations. Unfortunately, this essentially theoretical proposal has shown many practical limitations that we detail in Sup. Mat. A.1, notably important convergence issues, which explains its scarce use on real proteomics data.

More broadly, proteomic MV imputation aims at filling gaps in otherwise difficult to interpret biological data: So does multi-omic data integration. This field has elaborated on the natural assumption that a single omic modality contains only partial information which can be compensated for by applying multi-omics technologies to the same or related samples. To do so, one often estimates a common latent distribution modelling the underlying biological phenomena—classically using either matrix factorization [Argelaguet et al., 2018, Leppäaho et al., 2017, Rohart et al., 2017, Meng et al., 2019] or deep learning [Zhou et al., 2020, Barzine et al., 2020]). Then, the latent model is leveraged in a variety of tasks, such as pathway identification [Rohart et al., 2017, Argelaguet et al., 2018, Meng et al., 2019], sample extrapolation [Rohart et al., 2017, Barzine et al., 2020, Zhou et al., 2020], gene-set analysis [Meng et al., 2019], and possibly imputation [Flores et al., 2023]. It is worth noting that none of the available methods was specifically developed (and tested) to fulfil this task in the challenging context of bottom-up label-free LC-MS/MS data. We have however noticed a recent (still unpublished) attempt by Gupta et al. [2023] to train graph neural network in a gene-specific manner to impute protein-level MVs from such data, following a logic broadly akin to the one presented here.

In this work, we propose a concrete route to improve the imputation of LC-MS/MS-based label-free bottomup proteomic data by leveraging a well-established analytical and biochemical truism: precursors or peptides originating from the same protein should exhibit correlations that are insightful for proteomic MV imputation. Using this strategy, it becomes possible to accurately impute even with only few observed values per peptide, making our algorithm dramatically more performing than state of the art methods in the classical proteomic context of scarce samples. In addition, as insightful correlations can also be exploited between different omics modalities, we have generalized the approach to incorporate quantitative transcriptomic data, hereby opening the path to multi-omic based imputation of proteomic data. Doing so is however not sufficient to fully tackle the MNAR issue. Therefore, we have implemented these concepts using an imputation algorithm which estimates a single model for both censored and random MVs. Although our censorship model roots on that of PEMM [Chen et al., 2014], we propose a novel estimation strategy, more stable, scalable, and, unlike PEMM, without parameter-tuning or functional approximations, as to fit the practical constraints of proteomic data analysts. The resulting software, referred to as Pirat and released on the Bioconductor (https://www.bioconductor.org/packagesT7SS-ESX1-EccB-PF05108/release/bioc/html/Pirat.html), significantly outperforms all other imputation methods on a variety of tasks, datasets, and situations.

The article is organized as follows: The proposed method is described in Section 2. Then, sections 3 and 4 respectively present the validation protocol and results. Finally the paper ends with a discussion (Section 5) and conclusions (Section 6). All our results are reproducible with the code available on GitHub (https://github.com/TrEE-TIMC/Pirat_experiments).

## 2 Method

### 2.1 Notations

We hereafter use the following notations:

*X*: a complete matrix (of size *n*× *p*) of peptide log_2_ abundances, with rows referring to samples and columns to peptides. Note that depending on the context, *X* may refer to the entire dataset, or to a subset of it, namely a set of peptides or of precursor ions having the same mother protein(s).

- *X*_*i,j*_: the abundance value in *X* of the *i*-th row (sample *i*) and *j*-th column (peptide *j*); when *i* (respectively, *j*) is replaced by “.,” one refers to the to *j*-th column of *X* (respectively, the *i*-th column of *X*): *X*_.,*j*_ thus depicts the peptide vector (respectively, *X*_*i*,._ depicts the sample vector).
- *M* : an indicator matrix of the same size as *X*, reflecting whether an abundance value is missing or not. The same indexing notations as of *X* applies to *M* (*M*_*i,j*_, *M*_.,*j*_, *M*_*i*,._).
- *M*_*i,j*_: missingness indicator of abundance value of *j*-th peptide in *i*-th sample (1 if missing, 0 otherwise).
- *X*_obs_, *M*_obs_ (resp. *X*_mis_, *M*_mis_): all values of *X* and *M* corresponding to observed (resp. missing) abundance values.
- *x, m, etc*.: We use uppercase letters (*e*.*g*., *X*_*i,j*_ and *M*_*i,j*_) when considering random variables and lowercase ones (*e*.*g*., *x*_*i,j*_ and *m*_*i,j*_) when considering their realisations.
- *{i*, obs} (resp. {*i*, mis}): for a given sample *i*, set of (*i, j*) pairs such that *x*_*i,j*_ is observed (resp. missing). For clarity, brackets are removed when using this notation as subscript.
- *µ*: the vector of the theoretical mean abundance of the *p* peptides.
- Σ: the covariance matrix of the peptide’s abundances, of size *p*× *p*. The same indexing notation as for matrices *X* and *M* (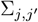 indicates the covariance between peptides *j* and *j*^*′*^).
- Γ: the set of parameters defining the missingness mechanism (see definition in Section 2.3).
- *I* denotes the identity matrix.

### 2.2 A bird’s eye view on Pirat methodology

Our method Pirat (standing for *Precursor or Peptide Imputation under Random Truncation*) works as follows: Firstly, depending on the input data, it creates either Peptide Groups or Precursor Groups. Both are based on protein belonging and are abbreviated as PGs, whereas peptides or precursors in a same PG are qualified as *siblings*. For sake of readability, we hereafter refer to peptides only, precursors being mentioned only if they require specific processing. It is usually assumed that sibling peptides will quantitatively behave similarly across treatment groups/biological conditions, even though the presence of isoforms, post-translational modifications, or miscleavage can impact these correlations [Dermit and Meyer, 2021]. Peptides from nested proteins form a unique PG and we duplicate peptides that are shared between PGs. The resulting biochemically informed dependence graph can optionally be enriched with transcriptomic data: for any PG, one then appends the sample-wise abundance vector(s) of related transcript(s).

Secondly, Pirat estimates over all the peptides a global missingness mechanism:

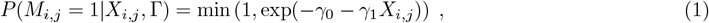

The missingness parameter Γ = (*γ*_0_, *γ*_1_) is estimated using a linera regression on a sliding window as to fit the missingness pattern of each dataset.

Thirdly, Pirat estimates the same model as PEMM [Chen et al., 2014], which is derived from the model of Little and Rubin [2019], yet using an original approach. Briefly, it aims at maximizing the penalized loglikelihood of observed data and missingness response, which, considering Γ known, amounts to maximizing the following quantity:

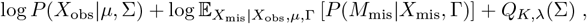

where *Q* is a penalty on Σ. In Pirat, we fit this model independently on each PG, with respect to *µ* and Σ. Likewise, penalty hyperparameters *K* and *λ* are automatically set for each PG in the empirical Bayes framework. To avoid the parametric approximations of the missingness mechanism that hamper PEMM’s results, we propose a tractable and differentiable lower bound of the above quantity. Therefore, the lower bound can be maximized with respect to *µ* and Σ using L-BFGS, a quasi-Newton optimization procedure proposed by Liu and Nocedal [1989]. To decrease computation costs and memory usage, as well as to enforce Σ positive definiteness, Pirat leverages an alternative parameterization of Σ based on its log-Cholesky factorization.

Fourthly, once the model is fitted, we propose to impute the MVs by their conditional mean with respect to observed values and their missingness response. The conditional mean being devoid of closed form, it is computed using standard Monte-Carlo integration [Robert and Casella, 1999]. The multiple imputed values obtained for shared peptides are averaged.

### 2.3 Model’s assumptions

Our model relies on the following four main assumptions regarding the whole peptide dataset (here, *X* and *M* refer to all the peptides of all the PGs of the dataset):

1. The *X*_*i*,._ are i.i.d. and normally distributed with parameters *µ* and Σ.
2. The missingness mechanism is self-masked for all peptides [Sportisse et al., 2020], *i*.*e*., the probability of an abundance value being missing, knowing the entire sample’s abundances, only depends on the abundance itself:

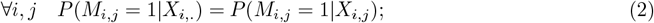 Or equivalently,

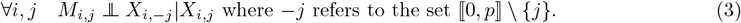
3. The missingness responses for each peptide of a sample are independent conditionally to their abundance [Sportisse et al., 2020]:

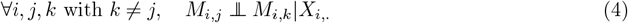
4. The missingness mechanism can be written as in [Chen et al., 2014] :

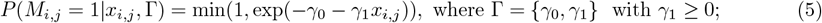 We assume here that there is a minimum detection threshold − *γ*_0_*/γ*_1_ below which no peptide can be quantified, and that the probability for a peptide to be missing exponentially decays with its log-abundance.

These assumptions ensure identifiability of *µ*, Σ, and Γ parameters in univariate and multivariate cases, as demonstrated by Sportisse et al. [2020] and by Miao et al. [2016].

### 2.4 Estimation of the missingness parameters

The set of parameters Γ over the whole peptide dataset is estimated once and for all as follows:

1. Sort the peptides by their observable mean, and denote by *α*_*i*_ the mean of the *i*-th peptide *i* in the ordered vector *α*;
2. For 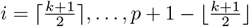 (where ⌈.⌉ and ⌊.⌋ denotes the upper and lower rounding), compute the rolling average of the missingness percentage *y*_*i*_ over the ordered peptides, with a window of size *k*; which leads to *p* − *k* + 1 points (*α*_*i*_, *y*_*i*_);
3. Fit a linear model on the points (*α*_*i*_, *y*_*i*_) by ordinary least squares, and set the parameters of the missingness mechanism (see Equation 5) as the coefficients obtained.

Using *k* = 10, preliminary experiments have shown a clear and smooth decreasing trend of log(*y*) with respect to *α* and a satisfying goodness of fit, as illustrated in Sup. Mat. Figure 11. Therefore, this value is used in all further experiments.

### 2.5 Penalized likelihood model

Considering the missingness parameters Γ known, we now propose to follow the selection model of Little and Rubin [2019] and to maximize the joint log-likelihood ℒ of the observed values and of the missingness response with respect to the Gaussian parameters *µ* and Σ, iteratively for each PG:

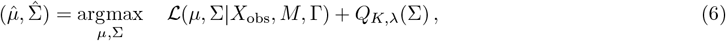

where *X* and *M* hereafter refer to the abundance matrix and missingness response of a single PG, and where *Q*_*K,λ*_(Σ) is a penalty term on Σ with hyperparameters *K* and *λ* (see Section 2.6), as proposed by Chen et al. [2014]. Hence, maximizing this penalized log-likelihood amounts to maximizing the posterior distribution of parameters *µ*, Σ, and Γ. Considering previous assumptions, the log-likelihood of the selection model decomposes as:

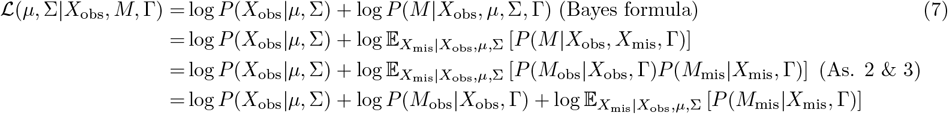

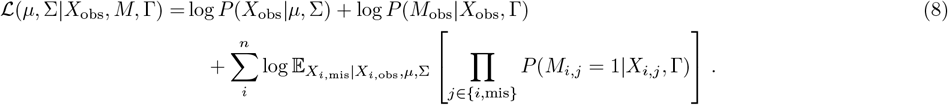

Note that the term log *P* (*M*_obs_ | *X*_obs_, Γ) does not depend on *µ* and Σ. Thus, if we consider Γ known, we can remove this term from the log-likelihood to optimize the parameters.

### 2.6 Penalty over Σ

We estimate Σ and *µ* for each PG independently by optimizing Equation 6. Therefore, the penalty hyperparameters *λ* and *K* must be tuned automatically (as many times as PGs). To do so, we rely on a Bayesian interpretation of the penalty term *Q*_*K,λ*_(Σ): it can be viewed as an inverse Wishart prior of the covariance matrix Σ. This penalty can be rewritten as:

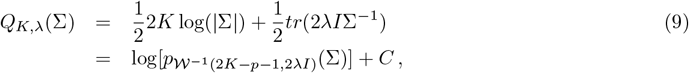

where 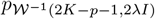 denotes the density of an inverse Wishart distribution with parameters (2*K*− *p*− 1, 2*λI*), and where *C* is constant with respect to Σ. Hence, *K* is related to the number of degrees of freedom in Σ estimate, and *λ* is related to the scaling matrix 2*λI*.

This leads us to empirically estimate the hyperparameters of each PG from the set of all fully observed peptides by relying on the empirical Bayes framework of Efron and Morris [1972] where we leverage that, in the univariate case, the inverse Wishart distribution boils down to the inverse-gamma one. Specifically, we compute their empirical variance and estimate the parameters of an inverse-gamma distribution (denoted *α* and *β*) through an MLE. Finally, by setting *K* = *α* + *p* and *λ* = *β*, the marginal prior distribution of each diagonal element of Σ (the variance of each peptide) is an inverse-gamma with parameters (*α, β*). Hence, the peptide’s variances have the same prior distribution among all PGs (this property is instrumental as before looking at any data, one would not expect the variance of a peptide to depend on the PG it belongs to). Without relaxing this property, we increase *α* by a factor 2 to better constrain the estimation of the covariance matrix (following Chen et al. [2014], which showed in various scenarios that increasing *K* improves the estimation of Σ).

### 2.7 Deriving a lower-bound of the penalized log-likelihood to estimate *µ* and Σ

A main issue regarding the estimation of *µ* and Σ by maximizing Equation 7, is that the expectation inside the log-likelihood is not analytically tractable for any missingness mechanism. The authors of PEMM proposed an approximation of the missingness mechanism that profoundly hinders the convergence of their model, as we show in Sup. Mat. A.1 with experiments on both real and synthetic data.

Another approach consists in using Jensen’s inequality, and the original missingness mechanism from Equation 5. We can then derive a tractable and differentiable lower bound of the likelihood that can be optimized. Concretely, to estimate *µ* and Σ for a given PG, we propose to maximize the following lower bound, ∀*j* ∈ {1,…*n*}

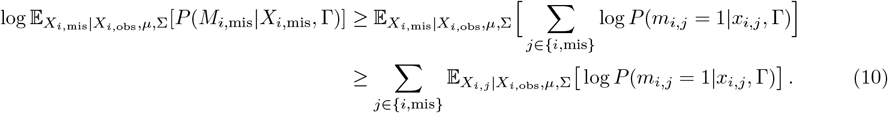

Let 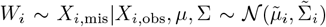 and *w*_*i*_ the realisation of *W*_*i*_. Using the parametric model from Equation 5, ∀*j* ∈ {*i*, mis}:

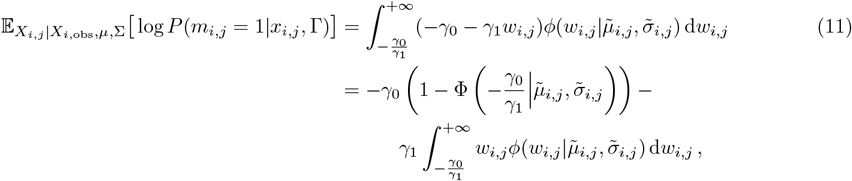

where *ϕ*(·|*µ, σ*) (resp. Φ(·| *µ, σ*)) denotes the (resp. cumulative) distribution function of a normal distribution with parameters *µ, σ*. Then, using properties of truncated normal distribution, we have:

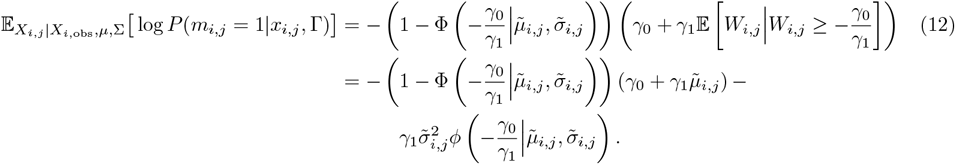

The expression of Equation 12 is differentiable with respect to parameters 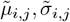.These two are the parameters of the conditional distribution of *X*_*i,j*_ with respect to *X*_*i*,obs_. They are obtained by the linear combination of *µ* and the Schur complement of the covariance matrix of observed values in *i*-th sample. Schur complement is differentiable with respect to Σ so that 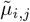 and 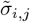 are differentiable with respect to Σ and *µ*. Hence, we can use any automatic differentiation tool, combined with optimization algorithm, to maximize the following lower bound of Equation 6:

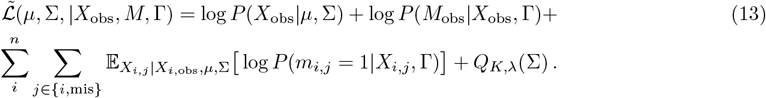

Specifically, we use L-BFGS (a quasi-newton method) combined with Armijo backtracking line-search, implemented with Pytorch [Shi and Mudigere, 2020].

To ensure Σ positive-definiteness during the optimization process, we re-parametrize Σ by its log-Cholesky factorization [Pinheiro and Bates, 1996]. Finally, to avoid ill-conditioned matrix that could cause numerical instabilities, we apply Tikhonov regularization method and subsequently add a fixed value *ϵ*_Σ_ = 10^−4^ to the diagonal of Σ. Doing so does not impact on the optimum solution, as the minimum variance observed in all the datasets processed so far has been ≥ 10^−3^.

### 2.8 Imputation

Once all parameters *µ*, Σ, and Γ are estimated, we impute missing values by their conditional mean. To do so, we use the following result of Chen et al. [2014]:

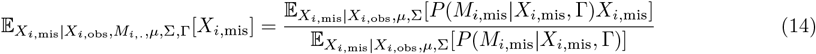

The distribution on which the left expectation is computed is not accessible, however, the distributions of the right expectations are Gaussian. Hence, we use Monte-Carlo integration to estimate them and compute the conditional mean of *X*_*i*,mis_. Note that, using the same approach, the order 2 moment of *X*_*i*,mis_ can be computed by simply squaring it in the above equation, which gives access the variance of *X*_*i*,mis_.

### 2.9 Imputation in singleton PGs

*Singleton PGs* (that correspond to proteins with a single and exclusive peptide or precursor ion being observed through MS analysis) regularly occur in proteomics datasets. Those PGs may be difficult to handle with Pirat, as no correlation can be leveraged. To cope with this we propose 3 altenatives: Pirat-2 (where singleton PGs are simply discarded), Pirat-S (standing for “Sample-correlations”) and Pirat-T (standing for “Transcriptome-informed”). Pirat-S and Pirat-T work as follows:

#### 2.9.1 Pirat-S

Pirat is based on the assumption the samples are independently distributed, see Section 2.3. Yet, other algorithms like ImpSec [Verboven et al., 2007] assume differently and leverage dependencies between samples. Following the same idea, we propose in Pirat-S to leverage sample-wise correlations to impute peptides of singleton PGs. To do so, we first estimate missingness parameters Γ and hyperparameters *K* and *λ* on the whole dataset and impute only PGs of size *>* 1 with the classical version of Pirat, described above. Second, we compute the empirical sample mean and sample-wise covariance matrix of the transposed imputed part of the dataset. Finally, we impute the remaining MVs by their conditional mean with respect to observed values, only using the observed values and the empirical sample mean and covariance matrix.

#### 2.9.2 Pirat-T

Pirat-T extends Pirat by enabling the integration of transcriptomic quantitative information, when available, as to guide peptide imputation. To do so, it requires a dataset of log_2_ mRNA expressions of samples from the same phenotype(s) as the proteomic ones. However, increments in imputation performances may require paired transcriptomic and proteomic samples. It also requires a correspondence table between transcripts and PGs, for instance, based on their original gene(s). In this work, transcriptomics is essentially used to improve singleton PG imputation (see Section 4.3). However, Pirat’s methodology is more versatile and the packaged code makes it possible to tune which PGs are imputed using complementary transcriptomic information, depending on their size.

Concretely, Pirat-T works as follows: for each PG, it integrates all the transcripts (*i*) for which non-zero values are measured in at least two conditions, and (*ii*) which are associated to at least one of the proteins of the PG. In the case proteomic and transcriptomic samples are paired, the log-count vectors of the transcripts are simply appended to the PGs. Otherwise, a condition-wise mean log-count vector is used instead. *In fine*, each mRNA log-count vector is processed like an additional (fully observed) peptide, and thus contributes to imputation depending on its correlation with the sibling peptide(s).

## 3 Validation protocol

### 3.1 Datasets

To conduct our experiments, we rely on the several publicly available datasets: Cox2014 [Cox et al., 2014], Bouysssie2020 [Bouyssié et al., 2020], Huang2020 [Huang et al., 2020], Capizzi2022 [Capizzi et al., 2022], Vilalllongue2022 [Vilallongue et al., 2022], Habowski2020 [Habowski et al., 2020], and Ropers2021 [Ropers et al., 2021]. A summarized description of how these datasets have been constituted is available in Supp. Mat. A.2. On all datasets, we remove peptides or precursors having strictly less than 2 observations. Finally, we apply log_2_ transformation both on peptide or precursor intensities and transcript normalized counts (when available).

### 3.2 List of competitive methods

To benchmark Pirat, we use 15 state-of-the-art imputation methods, many of which have already been pinpointed by NAguideR evaluation software [Wang et al., 2020]: Bayesian Principal Component Analysis (BPCA) [Oba et al., 2003] and Singular Value Decomposition (SVD) [Troyanskaya et al., 2001] parametrized with a number of components equal to the number of compared conditions in each dataset; GMS [Li et al., 2020] with 3 fold cross-validation for the tuning of its inner parameter (termed *λ*); K-nearest neighbors (KNN) [Troyanskaya et al., 2001] SeqKNN [Kim et al., 2004], trKNN [Shah et al., 2017] with the number of neighbors *k* set to 10; ImpSeq [Verboven et al., 2007] and its outlier oriented version ImpSeqRob [Branden and Verboven, 2009], Local Least Square (LLS) [Kim et al., 2005], MinProb [Lazar et al., 2016], MissForest [Stekhoven and Bühlmann, 2012], MLE [Love et al., 2014], msImpute in MAR and MNAR versions [Hediyeh-zadeh et al., 2023], and QRILC [Lazar et al., 2016] with default parameters. MissForest and LLS were tested with an input format where peptides are in columns and samples in rows, contrarily to all other methods. Amongst those methods, the following ones are specifically devoted to low abundance censored values: MinProb, msImpute mnar, QRILC, TrKNN.

While most methods require only 2 observed values (sometimes less) per peptide, GMS and trKNN require at least 3 of them, and both msImpute versions, 4 of them. This is an issue for two reasons: First, because, on controlled datasets, the biological variability is so small that not imputing those peptides before testing for differential abundance does not hamper the results, hereby artificially boosting the feature selection performances with respect to real-life cases. Second, because observing only two values is common for peptides nearby the detection limit, so that a concrete assessment on the MNAR imputation error is not possible with these algorithms. This is why, regardless of their performances on the biomarker selection task (see Section 4.1), GMS, trKNN, msImpute MNAR, and msImpute MAR could not be considered in the mask-and-impute experiments (see Section 3.3).

Likewise, three remaining state-of-the-art methods could not be included in the benchmark for various reasons: (*i*) Penalized Expectation Maximization for Missing Values (PEMM) [Chen et al., 2014], which had originally been validated on a simulated dataset with 100 proteins and could not scale up to an entire real-life peptide/precursor dataset as a result of its prohibitive computational cost; (*ii*) The PIMMS methods [Webel et al., 2023] which are based on deep-learning approaches, and which consequently require large cohorts (more than 50 samples) to be fully efficient; (*iii*) ProJect [Kong et al., 2023], a general purpose omics imputation method which code has not yet been commented, structured and packaged at the time of our evaluations, so that we could not make it work on the benchmark data. (*iv*) Few elder methods with previously demonstrated poor performances (*e*.*g*., zero imputation, mean imputation, accelerated failure time [Taylor et al., 2013], etc.) where not included in the benchmark, for sake of clear enough plots.

Finally, in our experiments, we did not consider any meta imputation algorithm: post-processor, like GSimp [Wei et al., 2018, Wang et al., 2022], ensemble imputation, like IMP4P [Giai Gianetto et al., 2020], chained imputation, like MICE [van Buuren and Groothuis-Oudshoorn, 2011], etc. The reasons are that their performances directly relates to the imputation algorithm they rely on, and that most of them can be extended to incorporate new input algorithms like Pirat.

### 3.3 Differential abundance task

On datasets with known ground-truth about differentially expressed proteins [Cox et al., 2014, Bouyssié et al., 2020, Huang et al., 2020], we compare our methods with those described in Section 3.2 using the following procedure: *(i)* Impute missing peptide (or precursor) abundance values. *(ii)* Test the all-mean equality of each peptide (or precursor) using the one-way ANOVA omnibus test available in the R package *stats* [R Core Team, 2013], and retain the resulting p-values. If a peptide cannot be imputed by a given algorithm (because of too few observed values), the test is performed nonetheless if the number of observed values allows for it, otherwise we set its p-value to one—see [R Core Team, 2013] for details about missing value restrictions. *(iii)* Display the precision-recall curves for the precision range [90%, 100%] (*i*.*e*., low FDP setting) for each method, and compute the global area under the curve, (*i*.*e*., on the precision range [0%, 100%]).

### 3.4 Mask-and-impute experiments

To precisely evaluate imputation errors, it has become customary to add pseudo-missing values with different MCAR/MNAR proportions, and to compute Root Mean Square Error (RMSE) or Mean Absolute Error (MAE) between imputed and ground-truth values, which read as:

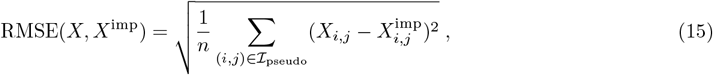

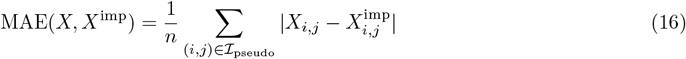

where ℐ_pseudo_ is the set of indices of pseudo-missing values and where *X*^imp^ is the imputed matrix. We generate pseudo MNAR values using a Probit left-censoring mechanism, *i*.*e*., the probability for a value *x* to be missing is 1 Φ(*x* | *ν, τ*), where the *ν* and *τ* are mean and standard deviation from the Gaussian cumulative distribution function Φ [Miao et al., 2016]. The overall missing value rate *α* and the MNAR/MCAR proportion *β* are controlled by applying the following procedure, adapted from Lazar et al. [2016]:

1. Set *τ* = *σ/*2 where *σ* is the overall standard deviation of the dataset.
2. For each value of *ν*^*′*^ on a linear scale between *µ* − 3*σ* and *µ* (the overall mean of the dataset), compute the expected overall rates *q* of missing values when applying Probit left-censoring wih parameters *ν*^*′*^ and *τ*, and retain these values. In practice we compute 100 points in total.
3. Interpolate points (*ν*^*′*^, *q*) obtained.
4. Compute *ν* associated to *βα* (the desired overall MNAR rate) with interpolated curve. If *αβ* is not in the range of interpolated curve, choose larger scale for *ν*^*′*^ at step 2.
5. Add MNAR values using Probit left-censoring with parameters *ν* and *τ*.
6. Add MCAR values with probability^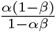.^

This method ensures that:

- The overall MNAR rate is *βα* (by construction of interpolated curve of (*ν*^*′*^, *q*)).
- The overall MV is rate is *α*, as

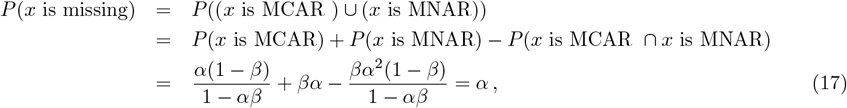

where *x* is an observed abundance value.

Of note, in the mask-and-impute experiments, all the methods are compared on the same artificially masked datasets.

## 4 Results

### 4.1 Pirat outperforms state-of-the-art on differential analysis task

We first evaluate our method on several biomarker selection tasks where significantly differentially expressed proteins are sought. We rely on three benchmark datasets for which differentially expressed proteins are known as their relative abundances are controlled, Bouyssie2020 [Bouyssié et al., 2020], Cox2014 [Cox et al., 2014], and Huang2020 [Huang et al., 2020]. They essentially involve a protein standard spiked in a complex yet constant biological background. These datasets have been produced with different MS acquisition modes (either data dependent or independent, a.k.a. DDA or DIA), at precursor- or peptide-level, and using different standard mixtures (see Supp. Mat. 3.1).

The feature selection is performed using the p-values resulting from the significance testing of the peptides’ differential abundance (described in Section 3.3), after imputing with our method (Pirat) as well as with 15 different state-of-the-art or popular methods (described in Section 3.2). Although receiving operating characteristic (ROC) curves are classically used to assess feature selection (they can be found in Sup. Mat. Figure 6), we have relied on precision-recall (PR) curves (Figure 1) instead. They provide broadly similar rankings, but the PR curves display the False Discovery Proportion (FDP), which directly relates to the FDR one often controls in proteomic experiments [Burger, 2018]. More precisely, FDR being classically controlled at 1% or 5%, we present on Figure 1 partial PR curves focused on the high precision (low FDP) region to better assess selection performances in this setting. Global area under the entire PR curve (AUCPR) are also given in Table 1.

**Table 1:** Global area under the precision-recall curves (and ranking into brackets) comparing Pirat with 15 imputation algorithms on a differential analysis task using three benchmark datasets (Bouyssié2020, Cox2014, Huang2020).

| Dataset | AUCPR % (rank) |  |  |
| --- | --- | --- | --- |
|  | A - Huang2020 | B - Cox2014 | C - Bouyssié2020 |
| BPCA | 67.26 (7) | 29.96 | 94.61 (3) |
| GMS | 65.38 (10) | 14.24 | 81.21 |
| ImpSeq | 67.20 (8) | 31.20 | 95.08 (2) |
| ImpSeqRob | 64.11 | 29.74 | 90.47 (9) |
| KNN | 59.33 | 51.94 (6) | 79.94 |
| LLS | SegFault | 31.48 (10) | 92.19 (8) |
| MinProb | 72.56 (3) | 81.08 (2) | 93.91 (6) |
| MissForest | 63.87 | 3.70 | 88.91 |
| MLE | 62.77 | 34.46 (8) | 23.58 |
| msImpute_mar | 71.78 (4) | 52.31 (5) | 94.22 (4) |
| msImpute_mnar | 73.88 (2) | 68.92 (4) | 94.14 (5) |
| Pirat | **77.02 (1)** | **86.43 (1)** | **97.22 (1)** |
| QRILC | 68.08 (5) | 80.76 (3) | 89.45 (10) |
| SeqKNN | 66.60 (9) | 34.91 (7) | 93.81 (7) |
| SVD | 67.32 (6) | 31.83 (9) | 89.03 |
| trKNN | 64.15 | 14.23 | 80.71 |

**Figure 1:**
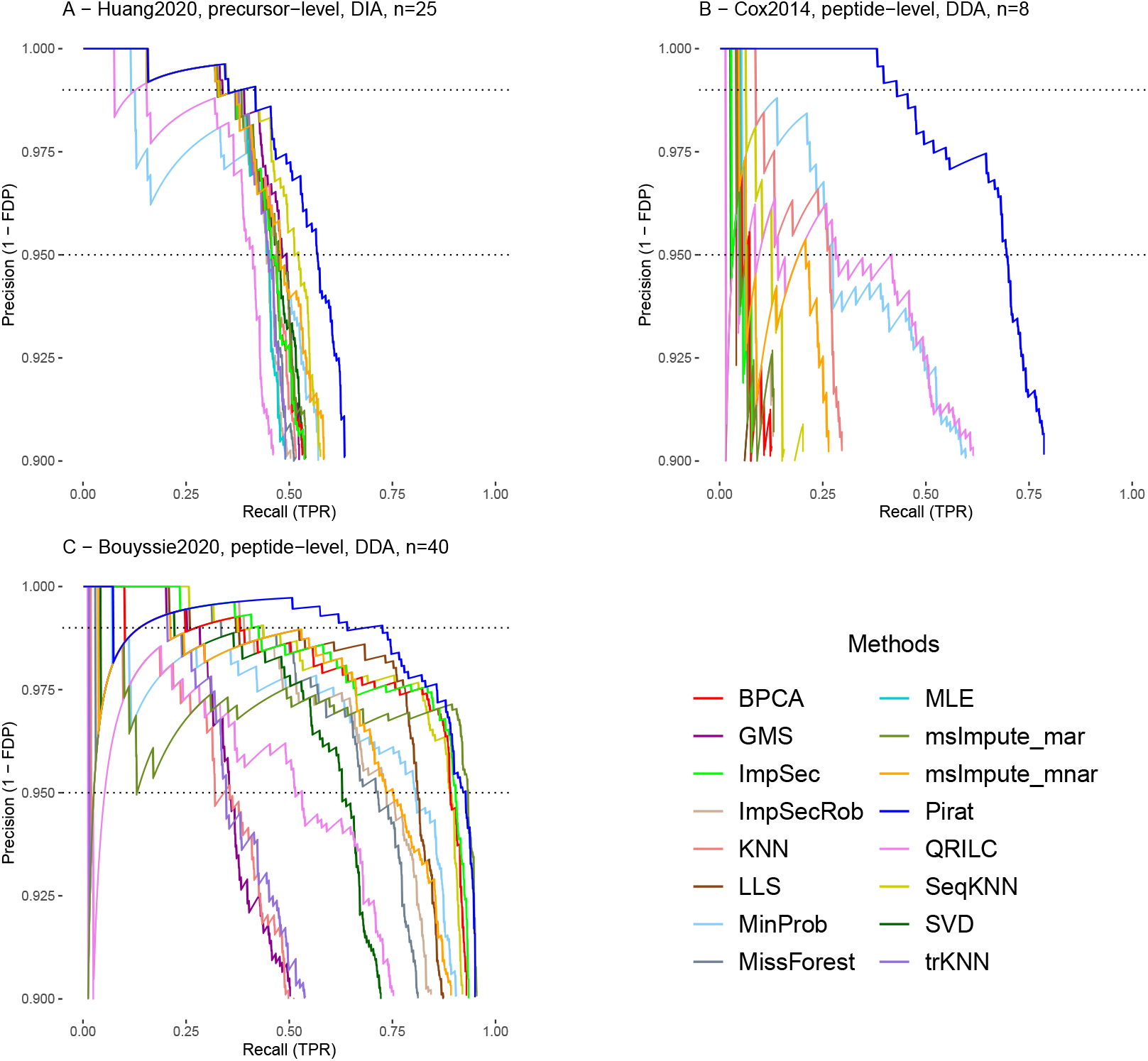
Differential abundance PR curves between 90% and 100% precision comparing Pirat to 15 imputation procedures on three benchmark datasets (A - Bouyssié2020, B - Cox2014, C - Huang2020), for which the name of the study, the imputation level (peptide or precursor), the type of acquisition (DIA or DDA), and the total number of replicates (*n*) are indicated in the subplot titles. The 1% and 5% FDP level are shown by the dotted lines.

On the three benchmark datasets, Pirat achieves the highest AUCPR (by a margin of 2.14% to 5.35% with respect to the second-best methods), thus indicating it best preserves the differential abundances while introducing less false positives. Precisely, Pirat provides the best PR trade-off on Huang2020 and Bouyssie2020 in the low FDP region, and almost entirely dominates the other methods on Cox2014. Beyond Pirat performances, these experiments are insightful with many respects: To begin with, several highly popular methods like MissForest, KNN or MLE have poor performances with respect to other less popular yet old methods like SeqKNN or ImpSec. Moreover, both msImpute methods (one of the most recent methods, published in April 2023) are very efficient, yet not as much as indicated in their seminal paper [Hediyeh-zadeh et al., 2023]. The reason is twofold: First, although well ranked in terms of global AUCPR, which concurs with the published results, they are outperformed by many other methods in the low FDP region (for example on Figure 1 A and C, SeqKNN exhibits a better PR tradeoff than both at low FDP). Second, they cannot impute missing values if less than four quantitative values are observed, so that the performances drop on datasets with fewer samples, like Cox2014. Conversely, Pirat clearly outperforms other methods on this dataset, which shows it can safely be applied in context of scarce samples. Most importantly, the lack of stability of other methods with respect to Pirat is noteworthy: Across the three datasets, the top scoring methods vary, first because the differences being sometimes marginal, the ranking is sensitive to random fluctuations; but also because of the changes in the MCAR/MNAR proportion. On Bouyssie2020, the MAR/MCAR oriented methods perform best after Pirat (ImpSeq, BPCA, and msImpute mar), whereas on Cox2014 and Huang2020, Pirat is followed by MNAR oriented methods (MinProb, msImpute mnar, QRILC). Overall, only Pirat remains in the top 3 for all datasets, and it does so with the first rank.

### 4.2 Pirat is robust with respect to its underpinning assumptions

Although Pirat displays the highest performances in an end-to-end task like biomarker discovery, understanding the extent to which the performance increments roots in the MS left-censoring or in the sibling peptide (peptides belonging to the same PG) correlations is insightful. To do so, we rely on mask-and-impute experiments (*i*.*e*., missing values are artificially added in real datasets, so that it is possible to measure the difference between the masked values and their imputed counterparts). To provide a baseline to the evaluations, we compare to the best-ranked methods according to the previous experiments. However, we have not been able to include msImpute mnar, msImpute mar, GMS, and trKNN irrespective of their performances, because of restriction on the number of observed values (see Section 3.2). Therefore, the subsequent evaluations compare Pirat with the best performing methods which can handle peptides with at least two observed values: MinProb, QRILC (MNAR scenario) and ImpSec, BPCA, SeqKNN (MAR/MCAR scenarios). Finally, a degenerated version of Pirat is also considered, where peptides are processed individually (*i*.*e*., in PGs or size 1), so that within-PG correlations cannot be leveraged.

In this experiment, we rely on datasets from two other LC-MS/MS proteomic analyses, detailed in Section 3.1. The first one, Capizzi2022 [Capizzi et al., 2022], is composed of 10 samples processed at the peptide level (16% of MVs). The second and third ones, Vilallongue2022 SCN and SC [Vilallongue et al., 2022], contain 8 samples processed at the precursor level (respectively 14% and 16% of MVs). According to our diagnosis tool (see Figure 2 second row), the sibling peptides tend to be strongly and positively correlated in Capizzi2022, as expected from a normal proteomic experiment. Oppositely, in Vilallongue2022 SCN, sibling peptides depict an odd correlation profile, and even more so in Vilallongue2022 SC, where the assumptions of bottom-up proteomics are hardly verified. Although questioning from an analytical viewpoint, these datasets are insightful, as they enable to evaluate PG’s benefits when siblings correlations are poor. For all datasets, we only consider the subset of peptides or precursors with no missing value, to which we introduce artificial MVs at a rate equal to that of the original dataset. Finally, we filter out peptides or precursors with one or zero remaining observed values. To evaluate the impact of the censoring model, which aims at improving MNAR imputation, these artificial MVs are a mix of MCARs and MNARs, where the former ones are generated according to a Probit mechanism [Miao et al., 2016], as described in Section 3.4. Note that this model used for masking is different from Pirat’s missingness model, which permits evaluating it in a misspecified setting. Then, we impute the missing values using the algorithms to benchmark and compute their associated Root Mean Square Error (RMSE) and Mean Absolute Error (MAE, given in Supp. Mat. A.5). We repeat this operation for different MNAR proportions (0%, 25%, 50%, 75%, and 100%) and seeds, leading to the curves of Figure 2.

**Figure 2:**
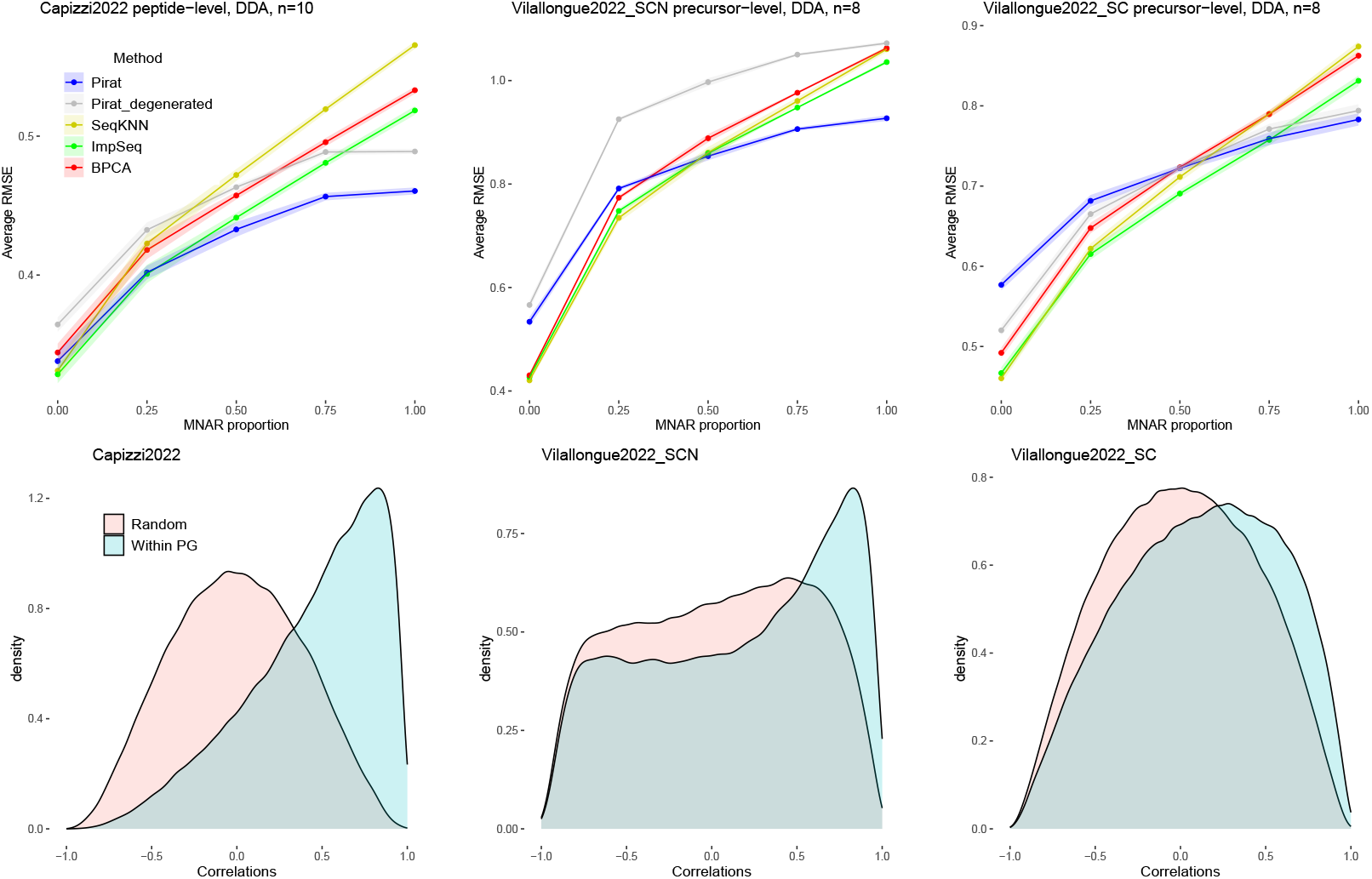
Average RMSE (top) of best imputation methods (according to the previous results) in function of the proportion of MNAR values on Capizzi2022 (left), Vilallongue2022 SCN (center), and Vilallongue2022 SC (right). The imputation level (peptide or precursor), the type of acquisition (DIA or DDA), and the total number of replicates (*n*) are indicated in the subplot titles. The errors averaged over 5 different seeds and margins correspond to standard deviations. On the bottom, empirical densities of correlations between peptides chosen randomly and between sibling peptides for each dataset respectively.

We do not display QRILC and MinProb results in Figure 2 as their MAE and RMSE is 3 to 4 times higher than that of other methods (full plots available in Sup. Mat. Figure 8). For those methods, the difference of performances between mask-and-impute and differential analysis is striking although easily explainable: these MNAR-devoted methods are highly biased towards low abundances and work in a univariate setting (*i*.*e*., apart from the knowledge of an ill-defined instrumental limit, no information is used), thus resulting in poor quantitative predictions. Yet, in cases where this low abundance is biologically relevant, erroneous predictions may not impede the accuracy of differential analysis: For instance, when a protein is not expressed in a phenotype, imputing its peptides with extremely low values may cause no significant harm.

All the other reference methods tested (PBCA, ImpSeq, and SeqKNN) have similar trends in terms of RMSE and MAE: As MCAR-devoted methods, the associated RMSE and MAE increase with the MNAR proportion. Although these three methods are broadly equally performing, ImpSec seems slightly but consistently more accurate, regardless of the MNAR ratio. As for Pirat, as expected, the performances vary according to the MNAR ratio and the quality of within-PG correlations, which validate the assumption at the root of Pirat algorithm. In more details, this variety of scenarios pinpoints the weaknesses of Pirat: in absence of MNAR values (which is not realistic in LC-MS/MS analyses), the performances are not as good because the instrument censoring model is useless. Likewise, in case of too low within-PG correlations, the performances decrease. However, when one out of the two assumption is fulfilled (significant MNAR ratio and poor within-PG correlations, or conversely, high within-PG correlations and large MCAR ratio) the performances are equivalent to the best state-of-the-art methods, both in terms of MAE and RMSE. More interestingly, as soon as both assumptions are fulfilled (which should be expected from most proteomics experiments), Pirat’s performances are unmatched, which explains why it has outperformed all other methods in the differential analysis task (see Section 3.3). Finally, we observe that Pirat outperforms in most cases its degenerated version. This is particularly showcased in MNAR dominant settings, which highlights the benefits of relying on PGs. On the opposite, it unveils the risk of lower-quality imputation for weakly-covered proteins.

### 4.3 Improving peptide imputation for proteins with low coverage

We now evaluate various options to best process the peptides of weakly-covered proteins. Precisely, we focus on singleton PGs, which are the most difficult situations for Pirat. As borderline cases, there is no unique satisfactory way to process them. The most straightforward and most statistically-grounded approach would be to adopt a quantitative version of the two-peptide rule often used at peptide identification [Munteanu et al., 2018], which would lead to filter out singleton PGs with missing values (see Supp. Mat. A.7). In case doing so is not acceptable for experimental reasons, we propose two alternatives: The first one (referred to as Pirat-S, see Section 2.9.1) is to leverage sample-wise correlations for imputation, with a similar logic as other algorithms (*e*.*g*., like ImpSeq or BPCA). The second one (Pirat-T, see Section 2.9.1) is to rely on complementary transcriptomic analyses, opening the path to a multi-omic view on imputation.

To evaluate them, we rely on two datasets involving both label-free proteomics and transcriptomics: Ropers2021 [Ropers et al., 2021] and Habowski2020 [Habowski et al., 2020]. Ropers et al. [2021] studied the effect of a synthetic growth switch on *E. coli* by performing paired proteomic and transcriptomic analyses of a wild-type and mutated strain at respectively 2 and 4 timepoints, thus resulting in 6 conditions (3 replicates each). In the mutated strain, protein and mRNA expressions are expected to decrease over time at different rate due to the differences of half-life of mRNA and proteins. As such, the samples have similar compositions leading to a relatively low MV rate at precursor-level (about 15%,) and we expect the transcriptomic-proteomic correlations to be exploitable thanks to the paired design.

In contrast, Habowski2020 results from unpaired transcriptomic and proteomic analyses of six types of mouse colon cells (3 replicates each for the proteomic experiments and 2 to 5 for the transcriptomic ones). Each condition being a different cell type, the sample composition in proteins varies considerably between the different conditions. This results in an important amount of precursor-level MVs (which reaches 50%) with values missing on an entire condition, a.k.a. MEC [Wieczorek et al., 2019]. Note that, compared to Ropers2021, this second dataset is crucial to (*i*) measure the possible negative impact of uncorrelated transcriptomic data in a multi-omic imputation approach; (*ii*) assess the importance of within-PG correlations to correctly impute poorly observed peptides; (*iii*) to evaluate if the performances are affected when samples are not perfectly correlated (leading to MEC-type MNARs). See Sup. Mat. 3.1 for more details on both datasets.

To evaluate the performance of these approaches, we rely on a mask-and-impute experiment similar to the previous one, with the additional goal to perturbate the true distribution of MVs the least amount possible. For this reason, we use the entire set of precursors (instead of using precursors with no MVs) and we add only a small amount of MVs (1% of the total number of true MVs in each dataset). This results in a few values that can be used to assess the performance of the approaches. We thus consider only the two extreme scenarios: 0% and 100% MNARs (respectively named MCAR and MNAR scenarios in the following experiments) and repeat this process ten times to assess the overall distribution of errors in each scenario.

#### 4.3.1 Pirat-S

Figure 3 shows, among others, the absolute errors of Pirat and Pirat-S. Clearly, Pirat-S improves upon Pirat in the MCAR setting for both datasets. This is not surprising: Pirat-S is conceptually close to ImpSeq, which has demonstrated excellent performances on MCARs (see Section 4.2). The improvement is more nuanced in the MNAR setting. Specifically, it seems to deteriorate the performances on Habowski2020. As pseudo-MNAR MVs are mostly carried by empty peptides (MEC patterns), too few values are left to derive informative correlations, notably when they do not originate from the same sample (see Sup. Mat. Figure 10, where the histogram E has a clearly different trend than histograms F, G and H). Although Pirat-S seems to be a safe choice in all other cases, medium-only imputation performances on MNARs with MEC (or similar) patterns reveal a limit of this approach.

**Figure 3:**
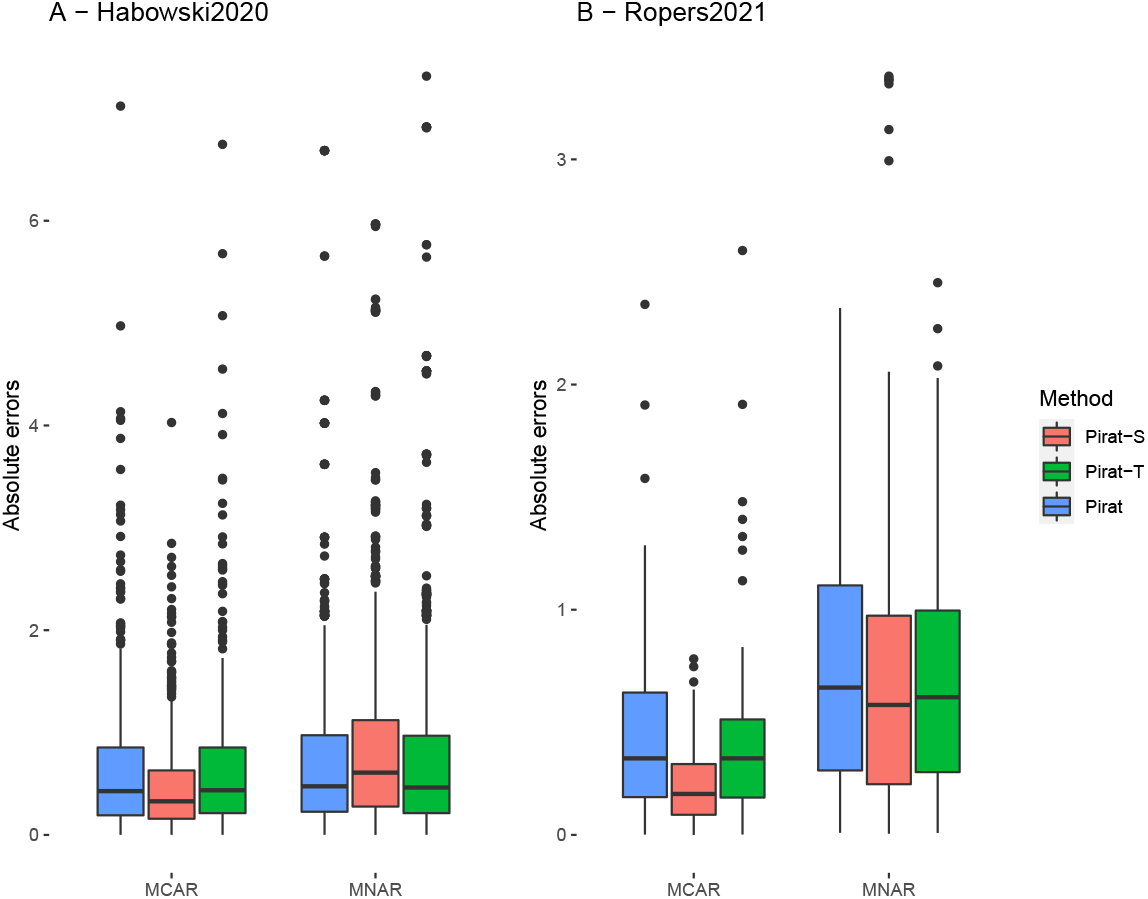
Boxplot of absolute errors for singleton PGs in A - Habowski202 and B - Ropers2021 datasets of Pirat, Pirat-S and Pirat-T.

#### 4.3.2 Pirat-T

Intuitively, the performance increment between Pirat and Pirat-T is expected to depend on the correlations between peptides and transcripts abundances. This is verified on Figure 3: the distribution error is shifted downwards on Ropers2021, where proteomic and transcriptomic samples are paired. On the contrary, the differences seem marginal on Habowski2020, where samples are not paired, thus making the transcriptomic patterns hardly exploitable on the proteomic data. Importantly, although transcriptomics is not helpful in this case, it does not deteriorate the performances either. This indicates that even in case a researcher wonders about the transcriptomic data quality or about their correlations with proteomics, it is not riskier to incorporate them for downstream imputation: Pirat-T robustly integrates transcriptomic data for imputation, without overinterpreting mRNA variations.

When comparing Pirat-S and Pirat-T, it appears the latter do not outperform the former, indicating that in general, sample-wise correlations can be more robustly leveraged than transcriptomic-proteomic ones (see Fortelny et al. [2017] for a discussion related to this topic). Although disappointing from a multi-omic integration perspective, this conclusion suffers one noteworthy exception: Habowski2020 on the MNAR setting; and interestingly, this is observed despite the lack of paired design. This suggests that in case of analysing proteomic samples from different tissues (with MEC-like MV values, which cannot be imputed accurately by leveraging sample or sibling peptide correlations) imputation can advantageously rely on transcriptomic assays. As those MEC values are essentially MNARs, it confirms the intuition shared in the proteomic community that complementary omic studies can be effective sources of information to explore more safely the proteome beyond the quantitation limit of mass spectrometry.

## 5 Discussion

Overall, Pirat outperforms previously published imputation methods across various experimental settings. On datasets endowed with differential abundance ground truth labels but with different missingness patterns and replicate numbers, Pirat consistently achieves the best PR trade-off, whereas other competitive methods display less stable and less accurate performances. Likewise, on mask-and-impute experiments, Pirat exhibits lower imputation errors for all scenarios with a majority of MNARs or with sufficiently good sibling peptide correlations. As for the other less realistic scenarios, the performances are more variable, yet close to the best ones in the worst cases. Indeed, while MCAR-devoted methods has long dominated the field of proteomics imputation, ignoring the physical limits of the instrument can only lead to biased results. Likewise, even though “discordant” peptides among siblings are likely to be observed as a result of unknown post-translational modifications or of poorly quantified peptides [Dermit and Meyer, 2021], they should be too scarce to affect the correlation distribution. Therefore, too low correlations should raise the experimenter’s awareness about the biochemical consistency of the assay. With this extent, the diagnosis tool accompanying Pirat is insightful. The above case suffers one exception, peptides without siblings (*i*.*e*., in singleton PGs), for which the difficulty to impute using Pirat does not relate to the experiment quality. To process them, we propose three alternatives to the original approach: (*i*) the quantitative two-peptide rule, *i*.*e*., discarding proteins with a single not fully quantified peptide. (*ii*) If different tissues are compared, it can be insightful to complement the assay using paired transcriptomics, as then Pirat-T can efficiently enhance the imputation of MNARs on singleton PGs. (*iii*) If transcriptomics is not available or not relevant for the proteomic study and if similar samples are analysed, sample-wise correlations can improve singleton peptide imputation using Pirat-S. With this regard, we draw attention to the marginal differences of global performances between the various extention of Pirat: Once averaged on an entire dataset, the imputation error on singleton PGs alone does not weight as much as the experimental design to drive the imputation strategy. We therefore encourage Pirat’s users to question their need with respect to proteins with low coverage before opting which strategy to use. Lastly, our experiments have reported Pirat’s excellent performances on peptide- and precursor-level datasets, thus ensuring its safe use in both cases. These results unfortunately do not allow us to formulate an educated opinion about which level is most appropriate to impute. Closing this debate would require testing the two approaches with a metric authorizing their comparison.

## 6 Conclusions

Considering its performances, we believe future research in proteomics imputation would advantageously leverage Pirat’s paradigms. Let us recall them: (*i*) the estimation of a global missingness mechanism inferred from the data; (*ii*) an explicit modeling of the biochemical dependencies that are known to result from the analytical pipeline; (*iii*) the possibility to include other omic measurements that are expected to correlate to the proteomic ones. Pushing further the developments in either of these three trends may require a different mathematical backbone than that of Pirat (*i*.*e*., not necessarily involving the penalized likelihood model of Equation 6), however, multiple paths are promising: regarding the missingness mechanism, a natural parameter estimator reads as the maximum likelihood estimate (MLE) over all the PGs. However, it requires replacing the optimization procedure described in Section 2.7 by a joint MLE, which memory and computation cost would be prohibitive. Fortunately, as Pirat processes each PG independently, it is highly parallelizable if acceleration is needed. Also, other missingness patterns like Probit or Logit [Li and Smyth, 2023] can fit the data, but before investigating them, some guarantees are necessary about their log-likelihood (or any lower bound, see Section 2.7) being tractable and differentiable. Besides, the self-masked assumption on the missingness mechanism only holds if we neglect the so-called “dynamic range” of mass spectrometers [Zubarev, 2013]. In practice, the variation of lower detection bound are not purely stochastic: they also result from which peptides are currently measured, notably the most abundant ones (as if highly abundant peptides could mask lower abundant ones). Thus, accounting for the intensity of co-eluting peptides in the missingness mechanism would be probably more accurate. A natural solution would be to add an extra term including relevant peptide abundances in Equation 5. However, this would require having access to additional data that are classically not reported, and extracting those pertaining to the dynamic range effect. From a biological standpoint, databases of known protein-protein interactions (PPI) are a source of supplementary correlation patterns, and incorporating them can be instrumental, notably for singleton PGs (as to find them pseudo-siblings capable of guiding the imputation). Likewise, known biologically or analytically relevant interactions can also be encoded in the prior of the covariance matrix Σ. For example, the correlations between protein-specific sibling peptides are expected to be larger than those involving either shared peptides, transcripts or PPI, so that it would make sense to adapt accordingly the scaling matrix prior described in Section 2.6. Finally, Pirat proposes to incorporate transcriptomic data, but other high-throughput technologies provide access to gene-wise or PG-wise quantitative information, which integration into Pirat follows the same logic (broadly speaking, one just appends the quantitative vectors of the various omics into a single matrix). Therefore, Pirat naturally opens the path to more elaborated integrative multi-omic methods to further tackle the challenge of proteomic data imputation.

## Author contributions statement

Funding acquisition (TB), research design and supervision (NV, TB), methodological development and prototyping (LE), data preparation (LE, LF, TB), experimental comparisons (LE), result analysis (LE, NV, TB), code and packaging (LE, SW), manuscript drafting (LE, NV, TB), manuscript assessment (LE, LF, SW, NV, TB).

## Acknowledgments

This work was supported by grants from the French National Research Agency: ProFI project (ANR-10INBS-08), GRAL CBH project (ANR-17-EURE-0003), SECRET project (ANR-22-CE45-0026), DEAP project (ANR-15-IDEX-02), and MIAI @ Grenoble Alpes (ANR-19-P3IA-0003).

## Availability of data and materials

The data and the methods are available on a publicly accessible Github repositories:

- https://github.com/prostarproteomics/Pirat
- https://github.com/TrEE-TIMC/Pirat_experiments

## A Supplementary Materials

### A.1 PEMM limitations and comparison with Pirat

We present in this section some experiments illustrating PEMM’s limitations. We rely on the original PEMM code, although the package is deprecated from the CRAN (as of 2022). First, we apply PEMM on all PGs of Ropers2021, with Γ parameters and *K* and *λ* hyperparameters obtained from the Pirat pipeline, as PEMM does not provide any way to tune them from the data. The other parameters are left as default. Among the 1830 PGs of the dataset, PEMM does not converge for 18% of them (a warning is returned in console), and the following error occurs in one PG: “Error in svd(X) : infinite or missing values in ‘x’”. In all non-converging cases, it happens because the likelihood becomes “NA” at some iteration. In addition, among these non-converging PGs, PEMM still returns 26% of the time a Σ matrix with a diagonal term exploding over 10^4^ (up until 10^50^). Moreover, a refined look at the behavior of the penalized likelihood computed by the PEMM function raises concern: On this dataset, it decreases in 16% of PGs. It does not necessarily happen when a PG does not converge; however, when it does, a Σ diagonal term explodes every time. We display on Figure 4 an example PG for which penalized likelihood decreases. Note that this behavior is in contradiction with the proof of convergence property of the penalized EM algorithm given in the original PEMM article. For all these reasons, PEMM is not included in Pirat’s benchmark.

**Figure 4:**
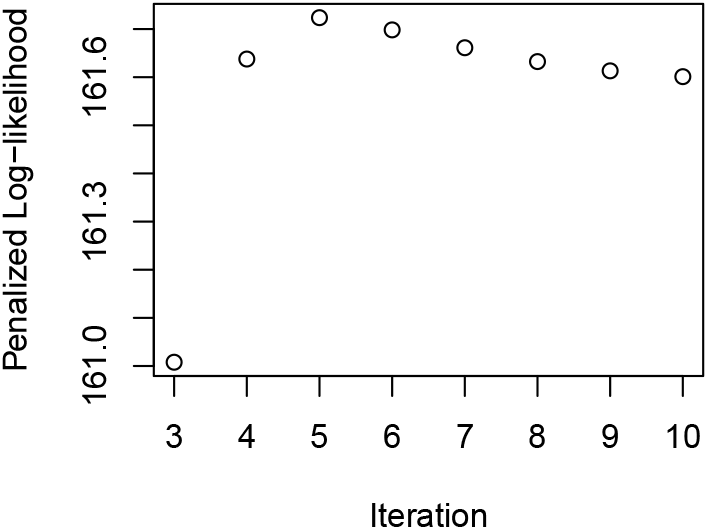
Penalized log-likelihood of PG 34 in Ropers2021 dataset, computed with `get.llk` function of PEMM package.

**Figure 5:**
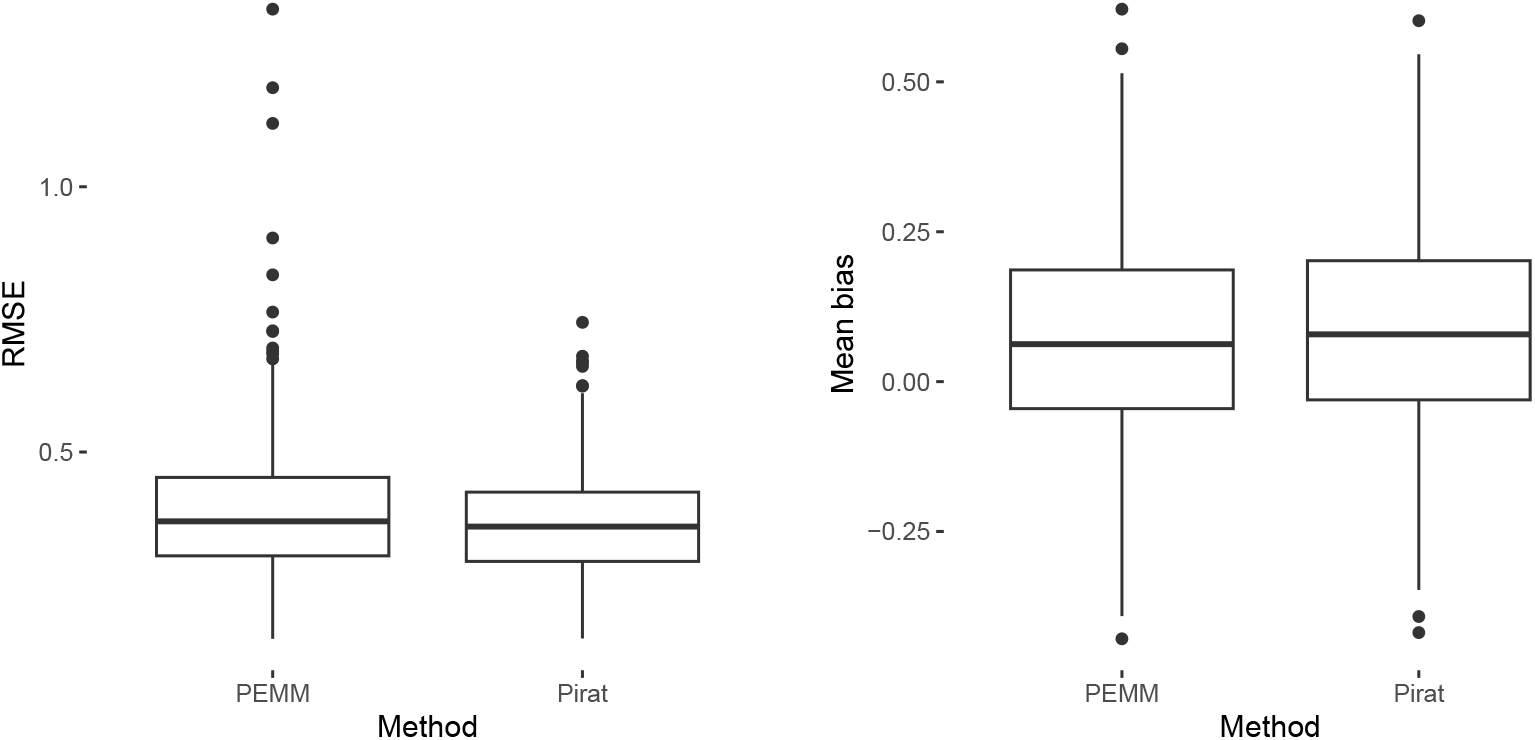
Boxplot of RMSEs between true and estimate *µ* among 1000 different synthetic datasets for PEMM and Pirat methods.

**Figure 6:**
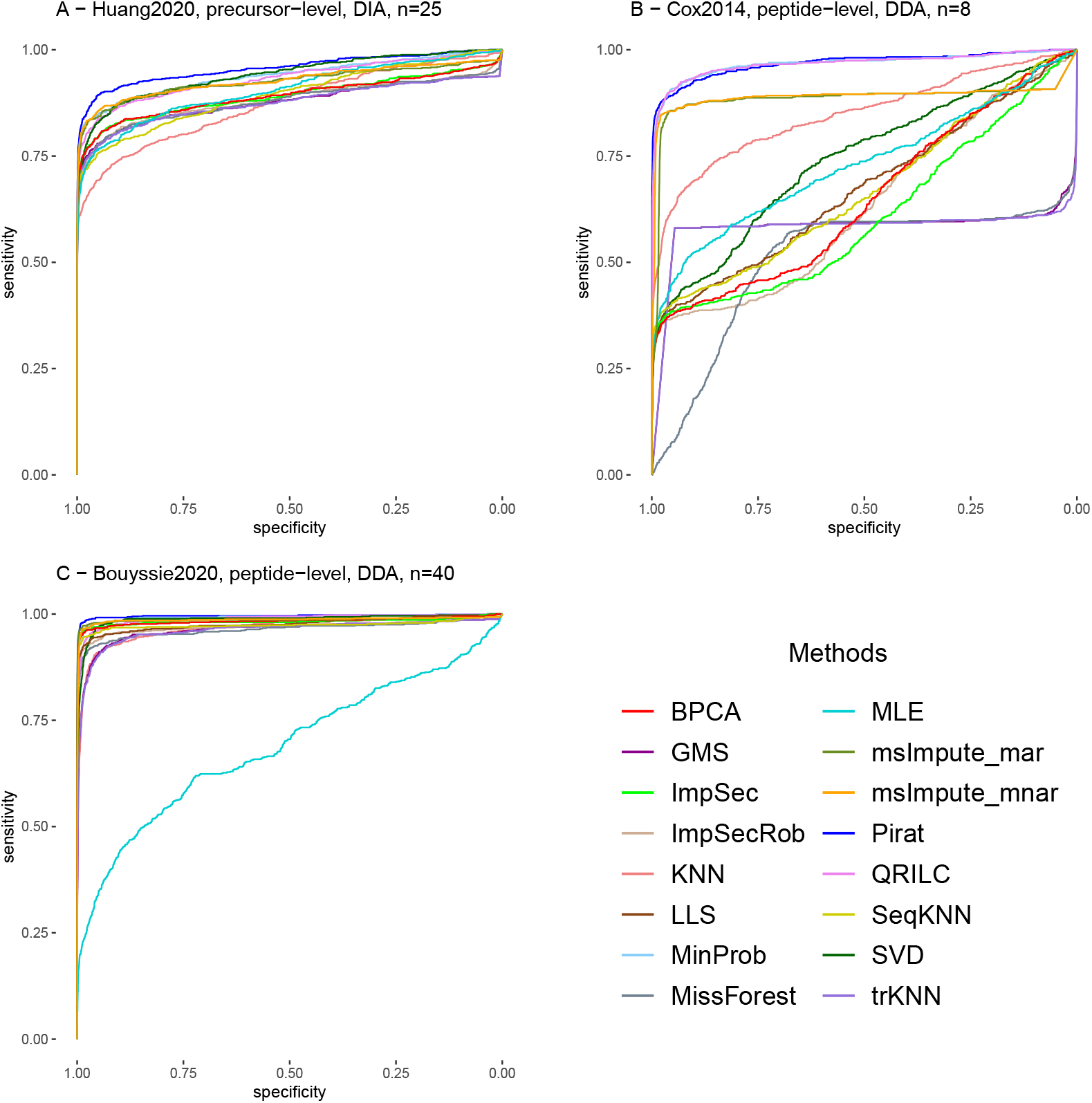
Complete ROC curves from p-values associated to the differential analysis validation on the three datasets Huang2020, Cox2014 and Bouyssié2020.

We claim that the convergence issue of PEMM relates directly to the analytical approximation used by the authors. In fact, a main issue regarding the estimation of *µ* and Σ by maximizing Equation 7, is that the expectation inside the log-likelihood is not analytically tractable for any missingness mechanism. It is why we choose in Pirat to derive its lower bound, which is provably tractable with PEMM’s missingness mechanism (see Section 2.7), instead of maximizing it directly. This expectation also appears in PEMM’s E-step. However, the author chose to compute it directly by approximating missingness mechanism Equation 5 as:

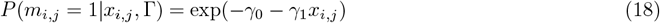

and by setting any observed value below − *γ*_0_*/γ*_1_ as missing. This approximation makes the log-likelihood unbounded and hinders convergence in some cases, according to the following argument. Using this approximation, we have (see Chen et al. [2014], Supp. Mat. Appendix D) ∀*i* ∈ {1, …, *n*}:

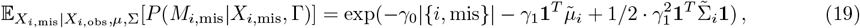

where 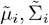 are the parameters of the normal distribution *X*_*i*,mis_|*X*_*i*,obs_. The above quantity diverges towards + ∞if any coefficient of 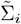 tends towards +∞, which is not consistent with the definition of a probability distribution. This analysis is consistent with the observed exploding variance terms during PEMM iterations in the previous experiment.

Beyond this essential improvement upon PEMM, Pirat differs on several other points that are listed in Table 2.

**Table 2:**
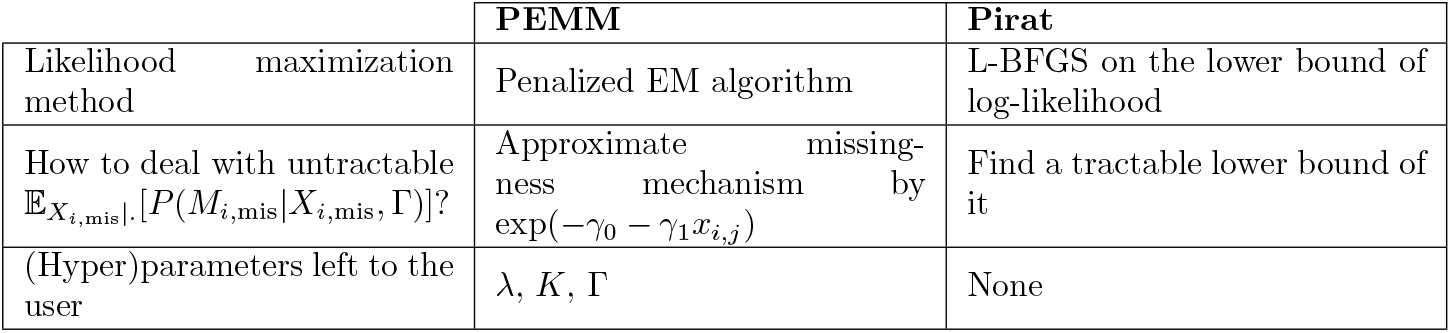
Summary of main methodological differences between PEMM and Pirat.

Despite the practical limitations of PEMM, it is useful as a reference for Pirat evaluations, to check if its estimation is not hindered by relying on the lower bound of the penalized log-likelihood. To this end, we compare the mean parameter *µ* mean bias and the RMSE on synthetic data, where the missingness mechanism is controlled, while being identical to the one used in Pirat and PEMM algorithms. We use 1000 simulated datasets generated with 10 features and 10 samples, from a multivariate Gaussian distribution with a random mean distributing alike Ropers2021 and a Toeplitz covariance matrix with exponential decay. We apply a mask on each dataset using the model of Equation 5, with parameter Γ chosen such that about 15% of values are missing (alike Ropers2021), and with *γ*_1_ *>* 0. We then apply PEMM and Pirat on each of them, providing both methods true Γ parameter. Finally, we compute the RMSE and mean bias between true and estimated feature mean for each dataset to subsequently display the corresponding box plots. As PEMM does not converge for 40% of the datasets generated (every time because of a “NA” likelihood), we only retain metrics of datasets for which PEMM converges (see 5). A Wilcoxon test between the distributions of the results of the two methods yields p-values of 0.03 (RMSE) and 0.12 (bias). To conclude, in this experimental setup where the missingness mechanism is known and provided and where the data leading to diverging estimates are ignored, Pirat estimate of *µ* is at least as good as that of PEMM.

### A.2 Dataset descriptions

We hereafter summarize the datasets used in our experiments, which are all publicly available, notably on ProteomeXchange. In parenthesis, we display for each of them the number of samples, peptides/precursors and PGs, respectively:

- **Cox2014** Cox et al. [2014] (8 samples, 22273 peptides, 2221 PGs): a mixture of 48 recombinant human proteins that are available as an equimolar mixture (UPS1) or mixed at defined ratios spanning 6 orders of magnitude (UPS2) with respect to UPS1 (10, 1, 10^−1^, 10^−2^, 10^−3^, 10^−4^). UPS1 and UPS2 are separately digested and resulting peptides are spiked into a *E. coli* lysate, resulting in two distinct experimental conditions. Each condition is analysed four times in data dependent acquisition (DDA, *i*.*e*., a single precursor is iteratively selected for fragmentation and identification by the mass spectrometer according to its response level in the first MS acquisition), resulting in four technical replicates per conditions. We use the file `peptide.txt` available at repository PXD000279, containing peptide-level abundances.
- **Bouyssie2020** Bouyssié et al. [2020] (40 samples, 20975 peptides, 2219 PGs): a mixture of UPS1 proteins spiked at different concentrations in the same yeast lysate. The dataset contains 10 conditions with 4 technical replicates each (each condition corresponding to a different UPS1 quantity, namely, 0.01-0.050.1-0.250-0.5-1-5-10-25-50 fmol, for 1*µ*g of yeast lysate). Hence, unlike Cox2014, within each condition, all the UPS proteins are spiked in with identical concentrations. Replicates are analysed in DDA mode. We use peptide intensities from `allsamples sum.xlsx` file available at repository PXD009815.
- **Huang2020** Huang et al. [2020] (25 samples, 132140 precursors, 6454 PGs): the UPS2 mixture spiked in tissue lysates from 25 mouse cerebellum samples. Five conditions (with 5 technical replicates each) are generated from spiked UPS2 proteins in known and different concentrations (namely 0.75-0.83-1.07-2.04-7.54 amol/*µ*l) into mouse cerebellum samples. The LC-MS/MS runs are acquired by the data independent method (DIA, *i*.*e*, a set of precursors in a given mass to charge range are iteratively selected for fragmentation and identification, in order to cover the whole MS acquisition, regardless of the measured intensities).

We use the published Spectronaut results in `Spike-in-biol-var-OT-SN-Report.txt` available at repository PXD016647, and imputed MS1 quantification at precursor level.

- **Capizzi2022** Capizzi et al. [2022] (10 samples, 14256 peptides, 2616 PGs): a study on Huntington’s disease effect on axonal growth in mouse. Two conditions with 5 biological replicates each are compared using LC-MS/MS in a data dependent acquisition mode (DDA), representing wild-type vs genetically modified model. Details of sample preparation can be found in the original article. The accession number for this dataset is PXD023885. Raw data is processed in the same manner as described in the original paper to obtain peptide-level quantification table.
- **Vilallongue2022** Vilallongue et al. [2022] (SCN:8 samples, 19050 precursors, 2741; SC:8 samples, 33364 precursors, 4238 PGs): a study on the influence of injury on visual targets in mouse. Five different tissue types are analysed by LC-MS/MS in DDA acquisition mode in independent quantification tables, with each time a control and an injured condition (4 biological replicates each). To test our method, we choose the two tissues that show the best and worse within-PG peptide correlations among the five (see 4.2): respectively, the suprachiasmatic nucleus tissue (SCN), and the superior colliculus (SC). The details of sample preparation can be found in the original article. The accession number for this dataset is PXD029325. Raw data is processed in the same manner as described in original paper to obtain precursor-level quantification table.
- **Habowski2020** Habowski et al. [2020] (18 samples, 26586 precursors, 3461 PGs): a study on the differentiation of mouse colon epithelial cells. Stem cells and 5 differentiated cells are analysed at proteomic and transcriptomic levels in an unpaired manner. Three biological replicates are used for each cell type in proteomic analyses. The transcriptome dataset is produced by Illumina paired-end stranded sequencing. The proteome dataset is produced by LC-MS/MS in DDA acquisition mode.
  - **Transcriptomic data processing** We download the raw sequencing data from GEO (GSE143915). Preprocessing and quality control are performed using the Trimmomatic Bolger et al. [2014] and FastQC tools respectively. Trimmed reads are aligned to the mouse GRCm39 genome assembly (filename given in Sup. Mat. Section A.3) by the STAR mapping software (version 2.7.8a) Dobin et al. [2013] provided with the Ensembl gene model given in Sup. Mat. Section A.3 for mapping splices. Read counts are generated using HTSeq (version 0.9.1; option -s no) Anders et al. [2015]. DESeq2 version 1.22.1 Love et al. [2014] is used to generate normalized counts.
  - **Proteomic data processing** We download the raw spectra from ProteomeXchange (accession ID: PXD019351). Peptides and proteins are identified using Mascot (version 2.7.0.1, Matrix Science) searching the Ensembl protein database for *Mus musculus* GRCm39 (filename given in Sup. Mat. Section A.3) appended with an in-house classical contaminant database. Mascot search is performed with the following parameters: trypsin/P as enzyme and two missed cleavages allowed; precursor and fragment mass error tolerances at 10 ppm and 20 ppm, respectively. The following peptide modifications are allowed during the search: Acetyl (Protein N-term, variable) and Oxidation (M, variable). The Proline software Bouyssié et al. [2020] is used to validate identifications: conservation of rank 1 peptides, peptide length ≥ 7 amino acids, false discovery rate (FDR) of peptide-spectrum-match identifications *<* 1% as calculated by Benjamini-Hochberg procedure and minimum of 1 specific peptide per identified protein group. Peptide ion intensities are calculated from extracted mass spectrum intensities of all peptides and normalized using variance stabilizing transformation with Prostar Wieczorek et al. [2017]. A more detailed description of the parameters used in the quantification step are provided in Supplementary Table 1.
- **Ropers2021** Ropers et al. [2021] (18 samples, 17472 precursors, 1830 PGs): a study on growth-arrested and wild type *E. coli* cells carrying a plasmid for glycerol production at various time points. A wild type and a modified strain are analysed in a paired manner at proteomic and transcriptomic levels. The modified strain is analysed at four different time points, resulting in four different conditions, and two others from the wild-type on the two first time points. Three biological replicates are made for each condition. The transcriptome dataset is produced by stranded sequencing on the Ion S5 using the Ion 540 chip. The proteome dataset is produced using LS-MS/MS in DDA acquisition model.
  - **Transcriptomic data processing**: Same procedure than for Habowski2020 is applied. GEO accession number is GSE168336. Trimmed reads are aligned to the reference *E. coli* K12 substrain MG1655 genome (Genbank assembly accession: GCA 000005845) provided with the Ensembl gene model given in Sup. Mat. Section A.3 for mapping splices.
  - **Proteomic data processing**:

The same procedure than for Habowski2020 is applied, except for the following: Raw spectra are downloaded from ProteomeXchange (accession ID: PXD024231). Peptides and proteins are identified by searching the same Ensembl protein database used in transcriptomic analysis (for this Ensembl genome assembly of *E. coli* the protein identifier is identical to the transcript identifier), appended with an in-house classical contaminant database, the plasmid pCL1920 protein sequences and the protein sequences of the GPD1 and GPP2 yeast genes which were cloned into it (Uniprot accessions: Q00055 and P40106). Carbamidomethyl (C, fixed) peptide modification is additionaly allowed during search. Peptides with length ≥ 6 amino acids are conserved at validation of identifications. A more detailed description of the parameters used in the quantification step are provided in Supplementary Table 2.

### A.3 Ensembl Gene models, genome assembly, protein databases

This section refers to the filenames of Ensembl gene model, genome assembly and protein databases used to build proteomic and transcriptomic datasets of Ropers2021 and Habowski2020 (see Section 3.1).

#### A.3.1 Ropers2021

The Ensembl genome assembly used for read alignment in transcriptomic analysis is the following: `GCA 000005845.2.fasta`

The Ensembl gene model used for mapping in transcriptomic analysis and for peptide identification in proteomic alignment is the following:

`Escherichia coli str k 12 substr mg1655 gca 000005845.ASM584v2.51.gtf`

#### A.3.2 Habowski2020

The Ensembl genome assembly used for read alignment in transcriptomic analysis is the following:

`Mus musculus.GRCm39.dna.primary assembly.fa`

The Ensembl gene model used for mapping slices in transcriptomic analysis is the following:

`Mus musculus.GRCm39.104.gtf`

The Ensembl protein database used for peptide identification in proteomic analysis is the following:

`Mus musculus.GRCm39.pep.all.fa`

### A.4 ROC curves

### A.5 Mean MAE for Capizzi2022 and Vilallongue2022 with best methods

### A.6 Mean MAE and RMSE with QRILC and MinProb

### A.7 Quantitative two-peptide rule (Pirat-2)

Even though it is technically possible, one should wonder whether it is reasonable to impute missing values for singleton PGs. First, protein abundances for those PGs will directly result from imputation. Consequently, the quality of the imputation will weight a lot in downstream analysis (*e*.*g*., differential analysis for biomarker discovery). Second, we cannot exploit the peptide-wise covariances for imputation, and thus suspect deterioration of the imputation for singleton PGs. To confirm this, we compare the median imputation error on singleton PGs with the error of PGs of size greater than two in the MNAR and MCAR settings. Results from this experiment (Figure 9) show that in most cases MVs from singleton PGs are indeed harder to impute than for all other PGs. Akin to the two-peptide rule in protein identification, a simple but effective strategy is to filter out singleton PGs that require missing value imputation to avoid errors in the imputation step to excessively affect downstream analysis. We call this strategy the *quantitative two-peptide rule* or simply the *two-peptide rule*.

**Figure 7:**
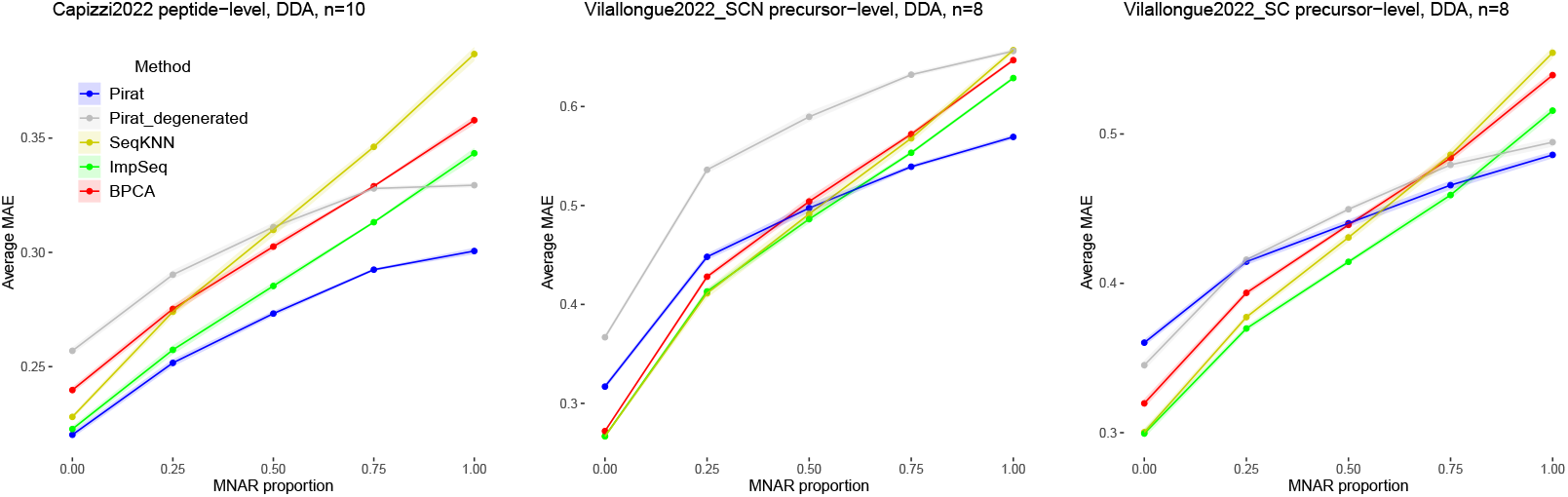
Average MAE (bottom) of BPCA, ImpSeq, Pirat, Pirat-Degenerated and SeqKNN in function of the proportion of MNAR values on Capizzi2022 (left), Villalongue2022 SCN (center) and Villalongue2022 SC. The imputation level (peptide or precursor), the type of acquisition (DIA or DDA), and the total number of replicates (*n*) are also indicated. The average MAE was computed over 5 different seeds, and margins correspond to standard deviation.

**Figure 8:**
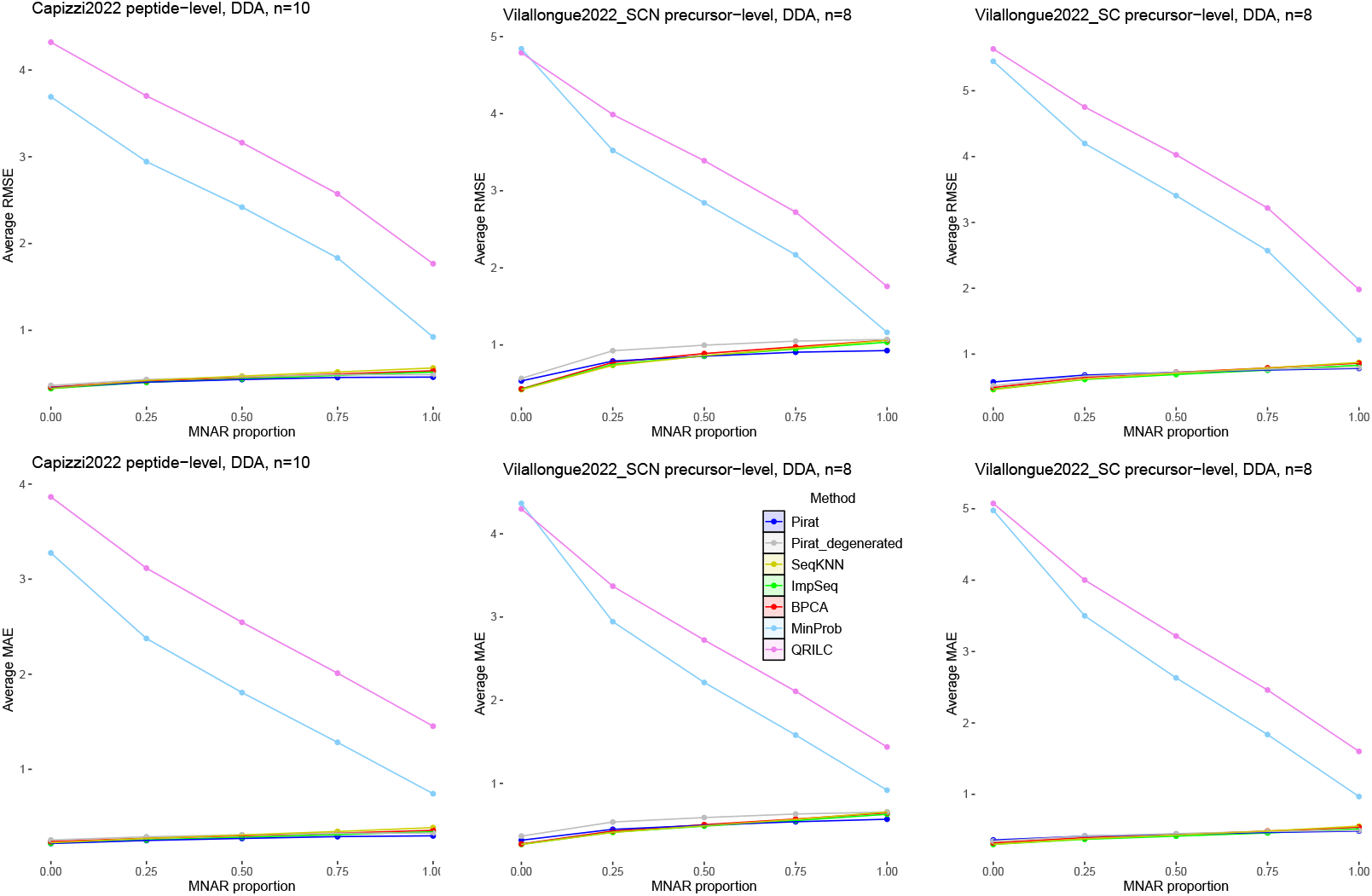
Average RMSE (top) and MAE (bottom) of MinProb, QRILC, BPCA, ImpSeq, Pirat, Pirat-Degenerated and SeqKNN in function of the proportion of MNAR values on Capizzi2022 (left), Villalongue2022 SCN (center) and Villalongue2022 SC. The imputation level (peptide or precursor), the type of acquisition (DIA or DDA), and the total number of replicates (*n*) are also indicated. The average RMSE and MAE was computed over 5 different seeds, and margins correspond to standard deviation.

**Figure 9:**
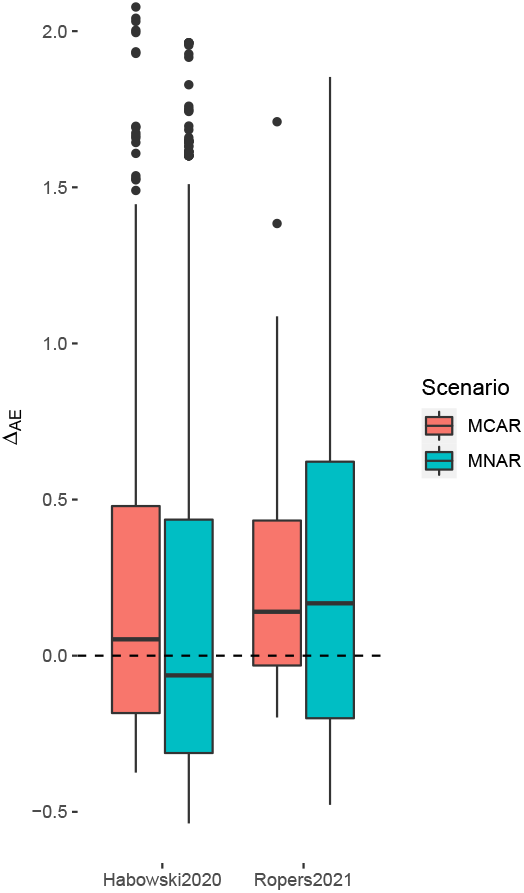
Boxplots of the distributions of the differences (denoted as Δ_*AE*_) between (*i*) the absolute errors in singleton PGs and (*ii*) the median of the absolute errors in all other PGs, after Pirat imputation, for a given dataset (Habowski202 or Ropers2021) and MNAR / MCAR scenario. Hence, for a given dataset and MNAR setting, the part of the boxplot that is above zero corresponds to absolute errors in singleton PGs that are greater than the median of absolute errors for non-singleton PGs.

**Figure 10:**
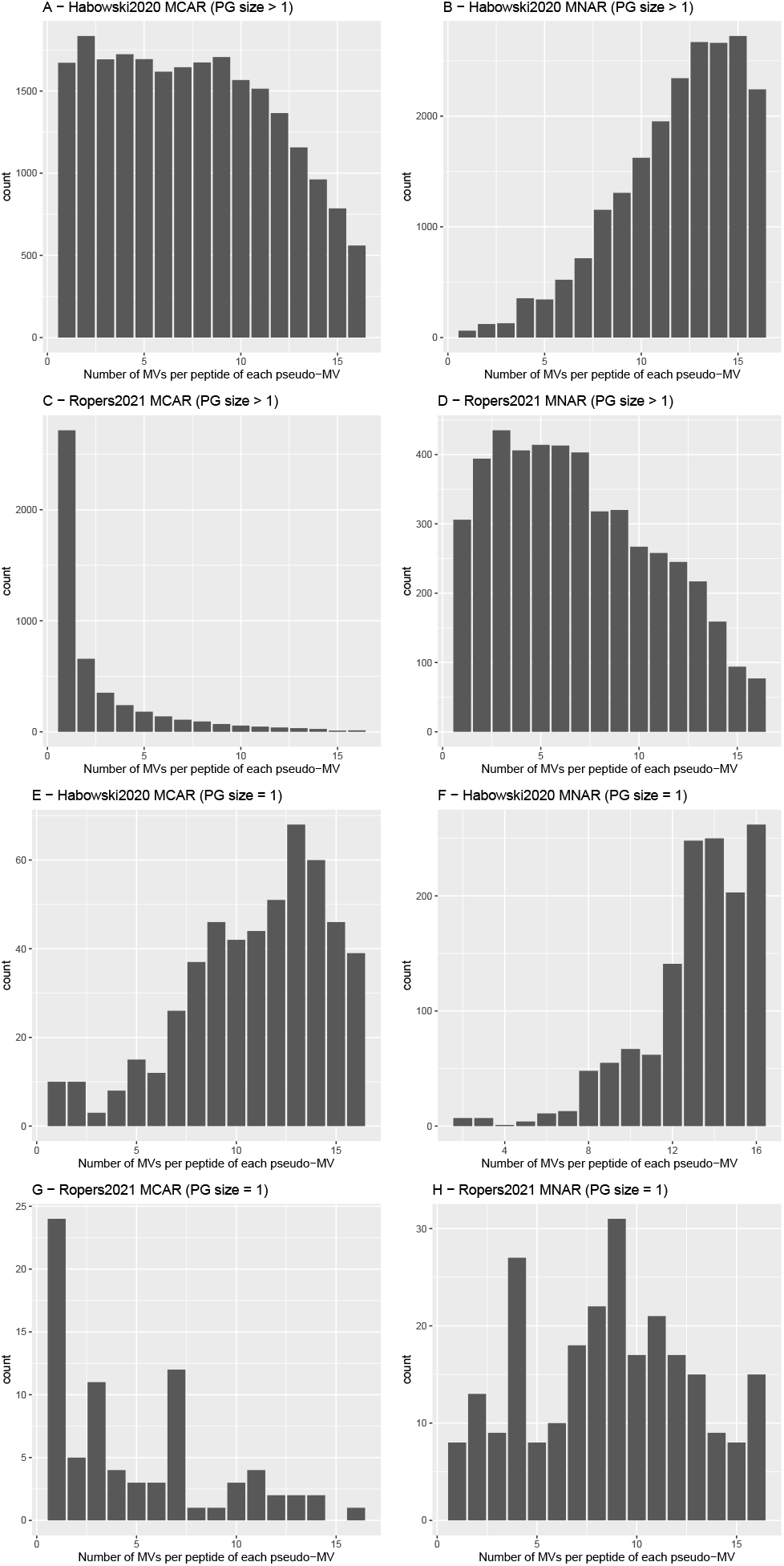
Histograms counting, for each pseudo-MV, the number of MVs (real or pseudo) in the same peptide, for Habowski2020 (A, B, E, F) and Ropers2021 (C, D, G, H), in MCAR (A, C, E, G) and MNAR (B, D, F, H) setting, and over 10 different seeds. We separate histograms for peptides contained in singleton PG (A, B, C, D) and non-singleton PGs (E, F, G, H). Note that the number of samples equals 18 in both datasets.

**Figure 11:**
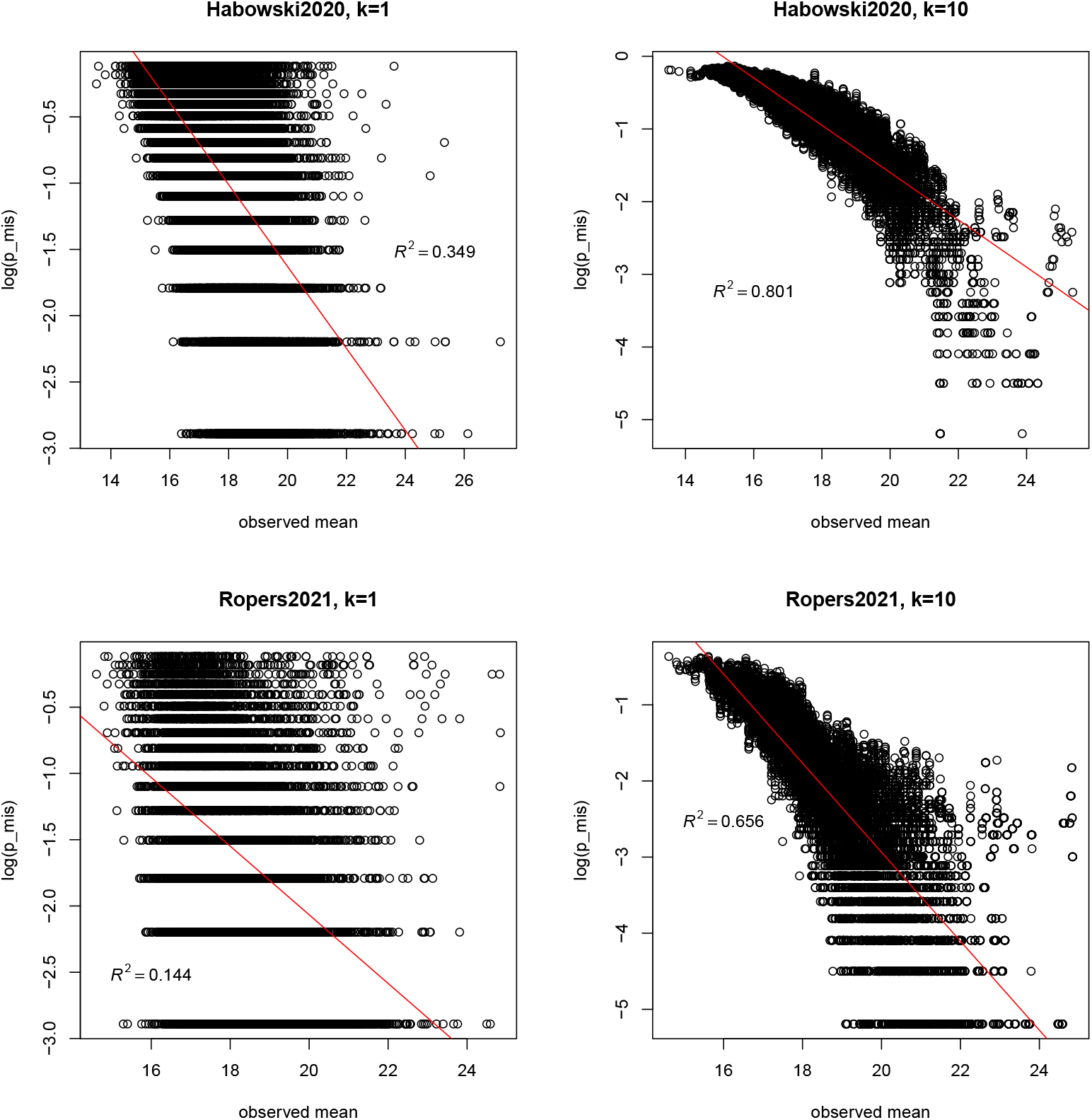
Regression of the log-probability of missing onto mean observed abundance following method described in 2.4, for Habowski2020 and Ropers2021, for *k* = 1 and *k* = 10. Residuals sum of squares (*R*^2^) of the linear regression are also displayed.

Looking more closely at Figure 9, the results of Habowski2020 in the MNAR setting are intriguing: we do not observe that the pseudo-MNAR values in this dataset are harder to impute on singleton PGs than on other PGs. A plausible explanation is that the pseudo-MVs of non-singleton peptides (which we compute the error on) in this setting are located on peptides containing mostly MVs. To confirm this, we show in Sup. Mat. Figure 10 (A, B, C, D) the histograms of the number of MVs contained in non-singleton-peptides carrying pseudo-MVs. We see that histogram A, representing Habowski2020 MNARs, has a clearly different trend from other scenarios: It is the only situation where pseudo-MVs are, for the vast majority, carried by very scarce peptides containing mostly MECs. Hence, correlations estimated by Pirat are uninformative as they are based on very few values coming from other conditions, which concurs with the drop of performances over non-singleton PGs with respect to singleton PGs in the MNAR setting.

### A.8 MV distribution in Habowski2020 and Ropers2021 experiments

### A.9 Fitting of missingness mechanism

